# Precision control of nanoparticle delivery with engineered biomimetic protein coronas

**DOI:** 10.64898/2025.12.09.693224

**Authors:** Tanveer Shaikh, Dhanush L. Amarasekara, Kenneth Hulugalla, Veeresh Torgall, Ryan J. Garrigues, Railey Mayatt, Thomas A. Werfel, Tonya N. Zeczycki, Nicholas C. Fitzkee

## Abstract

Nanoparticle delivery to tumors remains inefficient, with current nanomedicines achieving only 0.7% injected dose per gram (ID/g) of tumor tissue due to uncontrolled protein corona formation that redirects nanoparticles away from target sites. We engineered biomimetic protein coronas to control nanoparticle-protein interactions and enhance tumor targeting. Competitive binding studies using NMR spectroscopy revealed that transferrin (Tf) and fibronectin (Fn) outcompete albumin (BSA) and immunoglobulin G (IgG) for 15 nm gold nanoparticle surfaces, establishing a binding hierarchy that enables predictable corona composition. Pre-coating nanoparticles with a four-protein combination (BSA+Tf+Fn+IgG) created coronas that selectively enhanced cancer cell uptake while reducing macrophage recognition in vitro. When administered to tumor-bearing mice, these engineered coronas achieved 13 ppm/g tumor accumulation—equivalent to 4% ID/g—representing 6.5-fold improvement over bare nanoparticles and 2.6-fold improvement over PEGylated formulations. Proteomics analysis of secondary coronas formed in human serum revealed that engineered nanoparticles selectively recruit transport and adhesion proteins while limiting immune recognition signatures. The pre-formed coronas maintained targeting protein retention and reduced complement binding compared to controls. Circular dichroism confirmed minimal protein structural perturbation, preserving receptor-binding functionality for active targeting. The strategy harnesses natural protein adsorption processes to create “smart” biological interfaces that simultaneously evade immune clearance and promote tumor cell recognition through multiple receptor pathways. This approach demonstrates the feasibility of treating the coron as a programmable interface, addressing delivery limitations that have hindered clinical translation of cancer nanomedicines.

**For Table of Contents Only:** 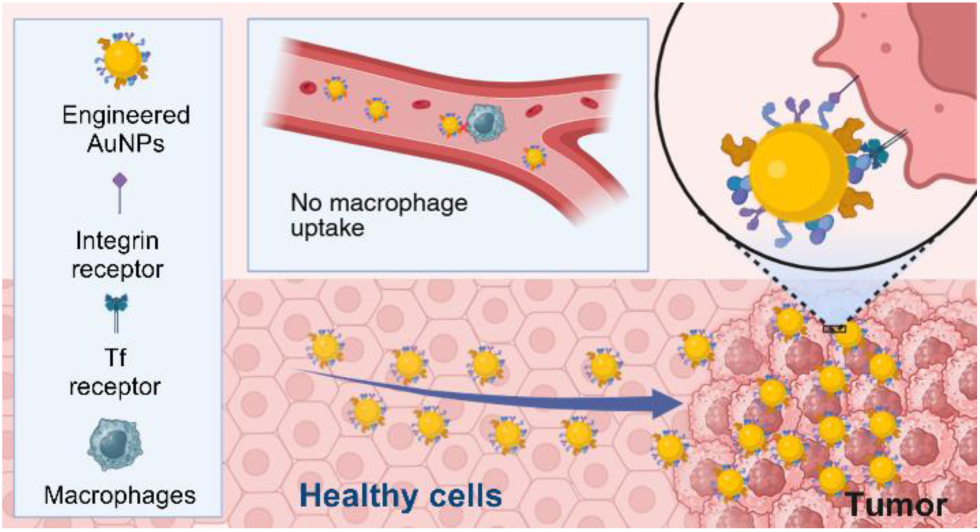

## Introduction

Nanoparticles (NPs) represent a promising platform for diverse biomedical applications, including drug delivery, bioimaging, and diagnostics. Their unique physicochemical properties, tunable surface chemistry, and ability to encapsulate therapeutic agents have positioned them as powerful tools in nanomedicine.^1, 2^ However, despite significant advances in nanoparticle design and functionalization strategies, their clinical translation remains challenging, with only a small fraction of administered nanoparticles reaching their intended target sites.^3^ A critical factor determining the biological fate of nanoparticles is the formation of a protein corona – a dynamic layer of biomolecules that adsorbs onto the nanoparticle surface upon exposure to biological fluids.^4, 5^ When nanoparticles encounter physiological environments, they rapidly acquire this complex protein coating that fundamentally alters their biological identity, influencing key parameters such as circulation time, biodistribution, cellular uptake, and therapeutic efficacy. The protein corona effectively creates a new biological interface that mediates interactions between the nanoparticle and biological systems, often overshadowing the intended design features of the original nanoparticle surface.^6–8^

Traditional approaches to modulate nanoparticle-biological interactions have focused primarily on surface modification strategies to prevent protein adsorption. Polyethylene glycol (PEG) coating, or “PEGylation,” has emerged as the gold standard for reducing opsonization and extending circulation time.^9–11^ However, PEGylation presents inherent limitations, including the “PEG dilemma,” where reduced recognition by the immune system comes at the cost of diminished cellular interactions, potentially compromising targeting efficiency.^12^ Furthermore, recent evidence indicates that even PEGylated nanoparticles develop protein coronas in vivo, albeit with altered composition and dynamics compared to unmodified counterparts.^13–16^

An alternative perspective has begun to emerge in recent years, viewing the protein corona not as an obstacle to overcome but as a biological feature that can be harnessed for therapeutic advantage. This paradigm shift has given rise to the concept of “protein corona engineering” – the rational design and manipulation of the protein layer surrounding nanoparticles to confer specific biological behaviors. By precisely controlling corona composition, structure, and dynamics, it may be possible to direct nanoparticle biodistribution, enhance cellular targeting, and improve therapeutic outcomes. The composition of the protein corona is influenced by multiple factors, including nanoparticle physicochemical properties (size, shape, surface charge), protein characteristics (concentration, affinity, molecular weight), and environmental conditions (temperature, pH, ionic strength).^17–23^ The corona formation follows complex kinetics, with high-affinity proteins initially binding to the nanoparticle surface (forming the hard corona), followed by the association of lower-affinity proteins (forming the soft corona). This time-dependent evolution of corona composition, known as the Vroman effect, adds another layer of complexity to predicting nanoparticle behavior in biological systems.^24, 25^

Recent studies have demonstrated that specific proteins within the corona can significantly influence nanoparticle-cell interactions. For instance, the presence of apolipoprotein E has been shown to facilitate nanoparticle transcytosis across the blood-brain barrier,^26, 27^ while clusterin has been associated with extended circulation time.^28^ Conversely, the adsorption of opsonins such as complement proteins and immunoglobulins can accelerate nanoparticle clearance by phagocytic cells.^29^ These observations suggest that selectively incorporating beneficial proteins while minimizing detrimental ones could potentially guide nanoparticle behavior in vivo. Despite these insights, several significant challenges remain in translating corona engineering into effective nanoparticle design strategies. First, the extreme complexity of biological fluids, containing thousands of different proteins at varying concentrations, makes it difficult to predict and control corona formation under physiological conditions. Second, the dynamic nature of the corona, with continuous protein exchange processes, complicates efforts to maintain a stable engineered corona throughout the nanoparticle’s journey in vivo. Third, the lack of standardized methods for corona characterization and quantification has hampered cross-comparison of results across different studies.^30, 31^

In this context, developing approaches to precisely engineer the protein corona with defined composition and structure represents a promising strategy to overcome these challenges. By pre-forming a corona with carefully selected proteins before exposure to biological fluids, it may be possible to create a more stable and predictable biological interface.^32–34^ This “biomimetic” approach, inspired by natural biological processes, could potentially combine the benefits of traditional surface modification strategies with the biological recognition capabilities of specific proteins. Several key proteins found in blood have attracted particular interest for corona engineering due to their abundance and known biological functions. Albumin, the most abundant serum protein, has been associated with extended circulation time and reduced non-specific cellular uptake.^35–37^ Transferrin, a glycoprotein responsible for iron transport, can facilitate receptor-mediated endocytosis in cells overexpressing transferrin receptors, including many cancer cell types. Fibronectin, an extracellular matrix protein, interacts with cell surface integrins and may enhance cellular adhesion and uptake. Immunoglobulins, while typically associated with opsonization and clearance, could potentially be utilized for targeting specific antigens when properly oriented in the corona.

Understanding the complex interplay between these proteins when co-adsorbed on nanoparticle surfaces is crucial for rational corona engineering. Competition between proteins for binding sites,^38^ conformational changes upon adsorption,^39^ and potential synergistic or antagonistic effects all influence the final corona composition and biological behavior. Developing methods to track individual proteins within complex mixtures and quantify their adsorption patterns represents a significant technical challenge that must be addressed to advance the field of corona engineering.

In this study, we present a systematic approach to engineer biomimetic protein coronas with precise control over composition and structure. Using 15 nm gold nanoparticles (AuNPs) as a model system, we investigate the competitive binding dynamics between key serum proteins – bovine serum albumin (BSA), transferrin (Tf), fibronectin (Fn), and immunoglobulin G (IgG) – and develop a strategy to track their behavior in complex mixtures using selective isotope labeling and NMR spectroscopy. By establishing a hierarchical binding model for these proteins, we rationally design engineered coronas with specific compositions that significantly enhance cancer cell targeting while reducing macrophage uptake in vitro. Moreover, we demonstrate that these engineered coronas substantially improve tumor accumulation in vivo, outperforming both conventional approaches like PEGylation and previously reported delivery efficiencies for nanomedicines. Our findings establish protein corona engineering as a powerful approach to modulate nanoparticle-biological interactions and enhance therapeutic delivery to target sites. By embracing rather than avoiding the protein corona, this strategy harnesses natural biological processes to achieve precision control over nanoparticle behavior in complex physiological environments. The principles and methodologies established in this work provide a foundation for rational design of next-generation nanomedicines with improved efficacy and translational potential.

## Results and Discussion

### Successful tracking of labeled BSA in a complex unlabeled protein corona

Our experimental investigations began by synthesizing 15 nm gold nanoparticles (AuNPs, **Fig. 1A**; Supporting Information, **Fig. S1, S2**) and incubating them with Bovine serum albumin (BSA) to confirm protein binding. Consistent with many prior studies, we observed a red shift in the localized surface plasmon resonance (LSPR) peak of the AuNPs upon incubation with proteins.^40^ This shift occurs because of a higher refractive index of the protein layer relative to water, which alters the dielectric environment around AuNPs, indicating protein corona formation. The LSPR peak shifted from 520 nm to approximately 525 nm when mixed with BSA (**Fig. 1B**). Dynamic light scattering (DLS) measurements revealed an increase in the hydrodynamic diameter of AuNPs upon incubation with proteins, with the size increasing from 17±3 nm to 45±4 nm when mixed with BSA (**Fig. 1C**). These findings conclusively indicate the formation of a protein corona when AuNPs are exposed to biological fluids. Our goal was to understand the complexity of protein corona behavior and how engineering different protein coronas affects the fate of nanoparticles, such as targeting ability, circulation time, etc. As a first step, we aimed to understand the competition between these proteins as they bind the AuNP surface. The ability to track individual protein behavior in the complex milieu of thousands of serum proteins was of particular interest.^5, 41, 42^ Our prior work used NMR spectroscopy to monitor the behavior of ^15^N-labeled proteins in unlabeled serum,^43^ but this approach is impractical for BSA, which cannot be easily isotope labeled.^44^ Instead, we attempted a protein modification approach with selective ^13^C labeling.

**Figure 1.**
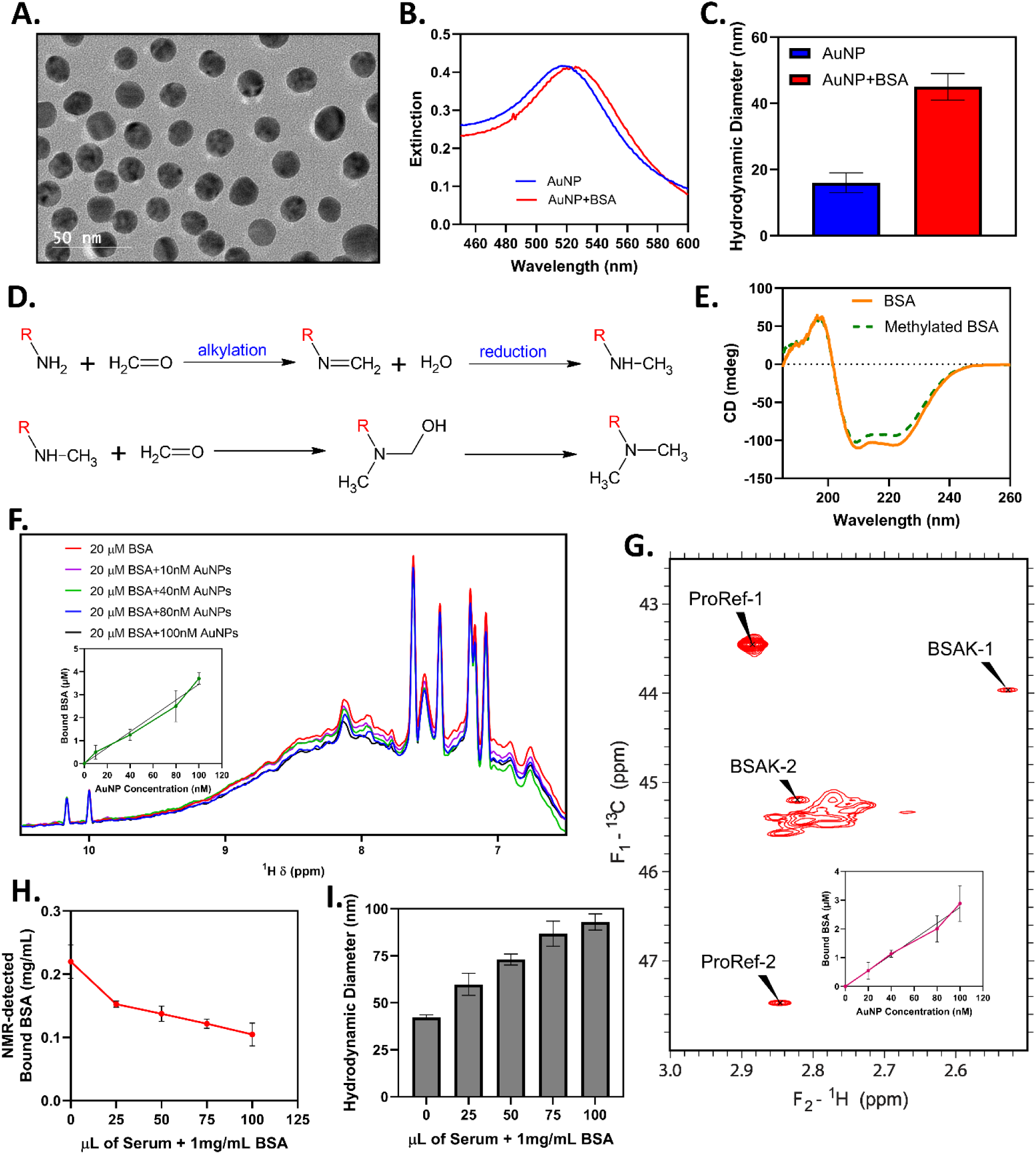
Development of a probe to track nanoparticle binding of an arbitrary protein in a complex mixture of proteins. (A) TEM images of synthesized 15nm AuNPs. (B, C) UV-Vis analysis and DLS analysis confirming the formation of protein corona. (D) Schematic reaction of ^13^C methylation using ^13^C-formaldehyde. (E) CD experiment to confirm no change in secondary structure of BSA after lysine methylation. (F, G) 1D ^1^H and 2D ^1^H-^13^C HSQC spectrum of BSA. The inset shows the plot of AuNP concentration vs Bound BSA to investigate the binding capacity. (H, I) NMR and DLS analysis showing the competition of serum proteins with BSA.

NMR spectroscopy offers distinct advantages over other techniques for quantifying protein corona information. It provides valuable structural insights, operates in a non-destructive manner, enables quantitative in-situ analysis, and exhibits versatility.^45–47^ In favorable cases, NMR can also be used to monitor the kinetics of nanoparticle adsorption directly.^48^ By using ^13^C methyl labeling, it is possible to monitor the behavior of ^13^C methyl groups in a complex mixture of unlabeled proteins.^49^ Formaldehyde is commonly used as a source of the ^13^C label, as it reacts specifically with the amine group of lysine residues in the presence of a weak reducing agent. Using ^13^C formaldehyde results in a stable N,N-dimethylated (^13^C) lysine residue (**Fig.1D**).^50^ The reaction is specific to lysine residues and does not significantly modify other amino acids in the protein. We began our investigation by examining the behavior of ^13^C-methylated BSA in the presence of AuNPs. In the past, we have used ^15^N-Trp as a quantitative external standard;^48^ here, for a ^13^C standard, we use 1 mM ^13^C-methylated proline, where the prolyl imine has been methylated and produces a strong, well-resolved set of signals in a ^1^H-^13^C HSQC spectrum (Supporting Information, **Fig. S3**).

In order to investigate the influence of methylation on the structure of BSA, we examined its circular dichroism (CD) spectrum. We observed no significant change in the CD spectrum of BSA following methylation, indicating that any alterations in the secondary structure were minimal (**Fig. 1E**). Next, we determined whether methylation affects nanoparticle binding capacity. Binding capacity (proteins bound per nanoparticle) can be determined for pure, unlabeled BSA using a simple 1D ^1^H NMR assay (**Fig. 1F**).^51^ The adsorption is calculated from the slope of intensity loss vs. AuNP concentration and was found to be 33±8 BSA per nanoparticle. The same nanoparticles were to determine the binding capacity of methylated BSA, this time using the ^13^C methyl signals in a 2D ^1^H-^13^C HSQC. While many signals are observed in BSA, three well-resolved signals were used for quantitation, two of which are shown (**Fig. 1G**). The adsorption capacity using this method was found to be 32±6 molecules of BSA per nanoparticle, in excellent agreement with the 1D proton experiment (**Fig. 1G**, inset). This confirms that both the 1D proton measurement and the 2D ^1^H-^13^C HSQC measurement yield consistent outcomes when assessing the adsorption capacity of BSA.

With an effective probe available for monitoring BSA in complex mixtures, we next wanted to see how serum influences BSA binding to AuNPs. Our 2D NMR experiments revealed that the amount of BSA bound to AuNPs decreased as the concentration of serum increased. Initially, 0.22±0.03 mg mL^−1^ of BSA out of 1 mg mL^−1^ BSA bound to AuNPs. However, with the introduction of serum into the system, this value decreased to 0.15±0.05 mg mL^−1^ and further declined to 0.10±0.02 mg mL^−1^ at higher serum concentrations (**Fig. 1H**). Clearly, other serum proteins compete with the BSA by effectively blocking the binding sites for BSA on the nanoparticle surface. Moreover, the corona of serum + BSA is more complex than BSA alone, as evidenced by the large increase in hydrodynamic diameter (94±5 nm) when serum is added to solutions of BSA. While this result supports the importance of competitive binding in the corona, we next sought to investigate specific competition between BSA and other abundant serum proteins found within the protein corona.

### Competition Dynamics Between Serum Proteins Reveal Strategies for Engineering Protein Coronas

It has been previously reported that other serum proteins can compete with BSA for binding to nanoparticles. For example, the serum protein apolipoprotein A-I (apoA-I) was found to compete with BSA for binding to gold nanoparticles in a study by Docter *et al.* using isothermal titration calorimetry.^52^ To investigate these competitive interactions systematically, we selected three abundant serum proteins – fibronectin (Fn), transferrin (Tf), and immunoglobulin G (IgG) – all commonly found in protein coronas.^53–55^ NMR experiments revealed distinct competition patterns for each protein. Both Fn and Tf significantly decrease the amount of BSA bound to the AuNPs in a concentration-dependent manner **(Fig. 2A, 2B)**, and DLS confirmed that they themselves become bound to the nanoparticle surface **(Fig. 2D, 2E)**. Interestingly, IgG exhibited markedly different behavior: it did not exhibit a concentration-dependent competition behavior (**Fig. 2C**). However, DLS of IgG and BSA mixtures exhibited substantial size increases, reflecting an IgG-mediated nanoparticle aggregation (**Fig. 2F**). This aggregation phenomenon aligns with observations by Chen *et al.*, who reported IgG-induced aggregation of gold nanoparticles in the presence of salt.^56^ Our results are consistent with previous studies, which have shown that other serum proteins can compete with BSA for binding to nanoparticles (e.g., fibrinogen) or can cause nanoparticle aggregation (e.g., immunoglobulins).^57–60^

**Figure 2.**
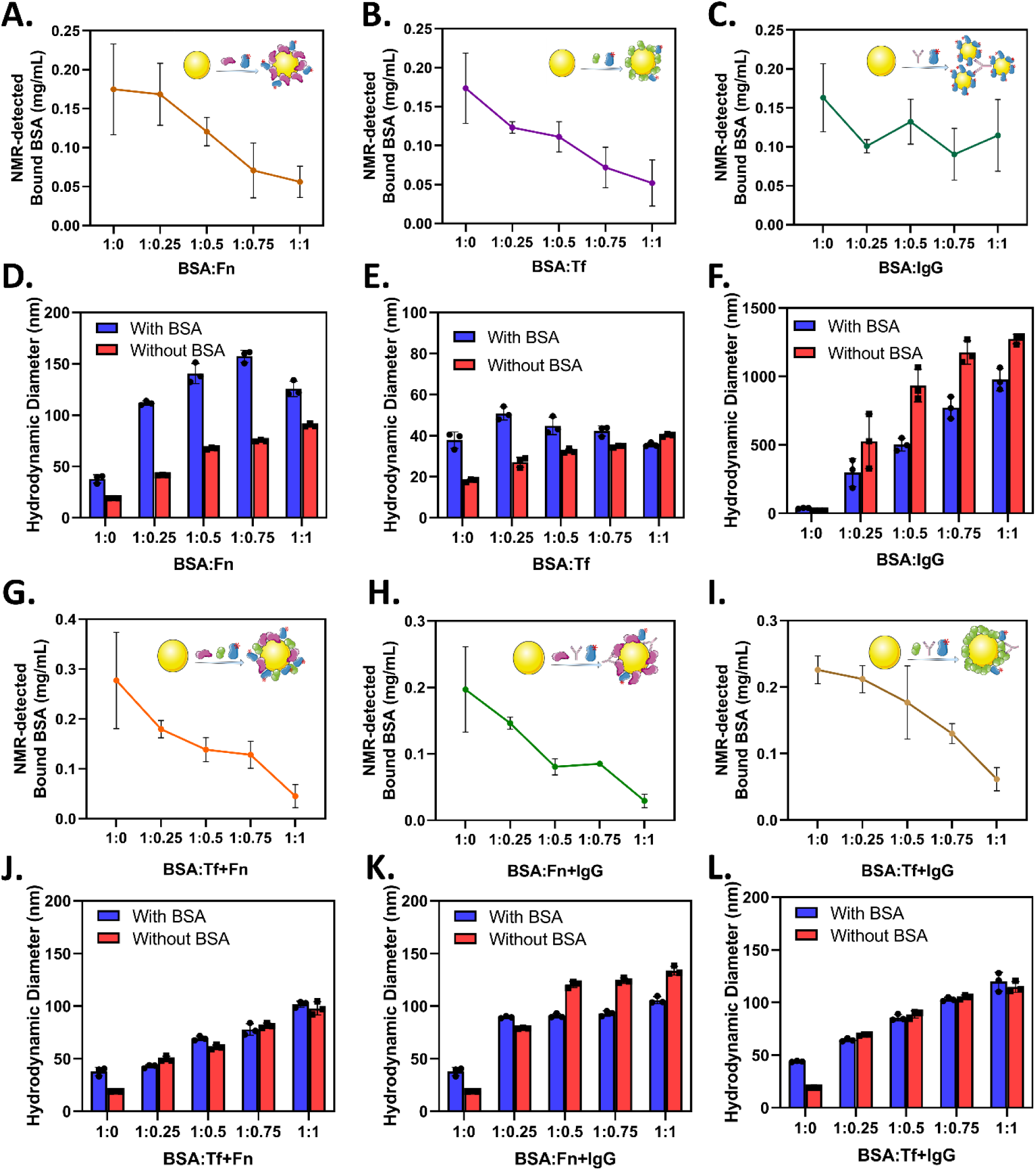
Competitive interactions between BSA and serum proteins on gold nanoparticles. NMR quantification of ^13^C-methylated BSA binding to AuNPs in the presence of increasing concentrations of (A) fibronectin (Fn), (B) transferrin (Tf), and (C) immunoglobulin G (IgG), shown as protein ratios (physiological serum ratio relative to 1 mg mL^−1^ BSA). Corresponding DLS measurements are shown for AuNPs incubated with (D) Fn, (E) Tf, and (F) IgG in the presence (blue) or absence (red) of BSA. NMR detection of BSA binding in three-component protein mixtures: (G) BSA:Tf:Fn, (H) BSA:Fn:IgG, and (I) BSA:Tf:IgG at varying protein ratios. Again, DLS measurements for these mixtures are shown: (J) BSA:Tf:Fn, (K) BSA:Fn:IgG, and (L) BSA:Tf:IgG. Error bars represent the standard error of the mean (SEM) of three independently prepared samples.

To better approximate physiological conditions while maintaining experimental control, we investigated three-component protein mixtures. NMR analysis revealed that all tested combinations, Fn+Tf, Fn+IgG, and Tf+IgG competed with BSA for binding to AuNPs (**Fig. 2G, 2H, 2I**). Notably, the competition patterns in these mixtures differed from those observed with individual proteins. DLS measurements of these three-component systems showed size increases larger than those seen with individual proteins, confirming the formation of complex, multi-protein coronas (**Fig. 2J, 2K, 2L**). However, a particularly striking finding was that IgG no longer caused aggregation when combined with either Fn or Tf, suggesting that these proteins modulate IgG’s interaction with the nanoparticle surface. When all four proteins (BSA, Fn, Tf, and IgG) were combined at concentration ratios approximating normal serum levels, the bound BSA was reduced to 0.09±0.02 mg mL^−1^ from an initial concentration of 1 mg mL^−1^. DLS indicates similar sizes of AuNPs at mixture compositions, indicating the saturation of proteins on the AuNPs surface (Supporting Information, **Fig. S4**). Despite the high concentrations of BSA and IgG in this mixture, they were effectively outcompeted by Tf and Fn, highlighting the hierarchical nature of protein binding affinities.

The competitive protein binding dynamics were further confirmed by zeta potential measurements, which revealed how different protein combinations alter the zeta potential of nanoparticles (Supporting Information, **Fig. S8**). For example, while IgG-coated nanoparticles exhibited a positive zeta potential, this became negative when Tf or Fn were present, providing additional evidence that these proteins successfully compete with IgG for the nanoparticle surface. These electrophoretic changes complement our NMR findings and demonstrate that the hierarchical protein binding we observed translates to measurable alterations in nanoparticle surface properties.

To determine the precise composition and quantification of these engineered coronas, we performed gel electrophoresis followed by densitometric analysis (**Fig. 3A**). The results revealed distinct specificity in protein adsorption patterns. When all four proteins were present in a concentration ratio reflecting what is seen in blood serum, Tf and Fn dominated the corona composition, despite the higher concentrations of BSA and IgG in the mixture. This preferential adsorption is clearly visualized in the heat map (**Fig. 3B**), which shows the relative abundance of each protein across different combinations.

**Figure 3.**
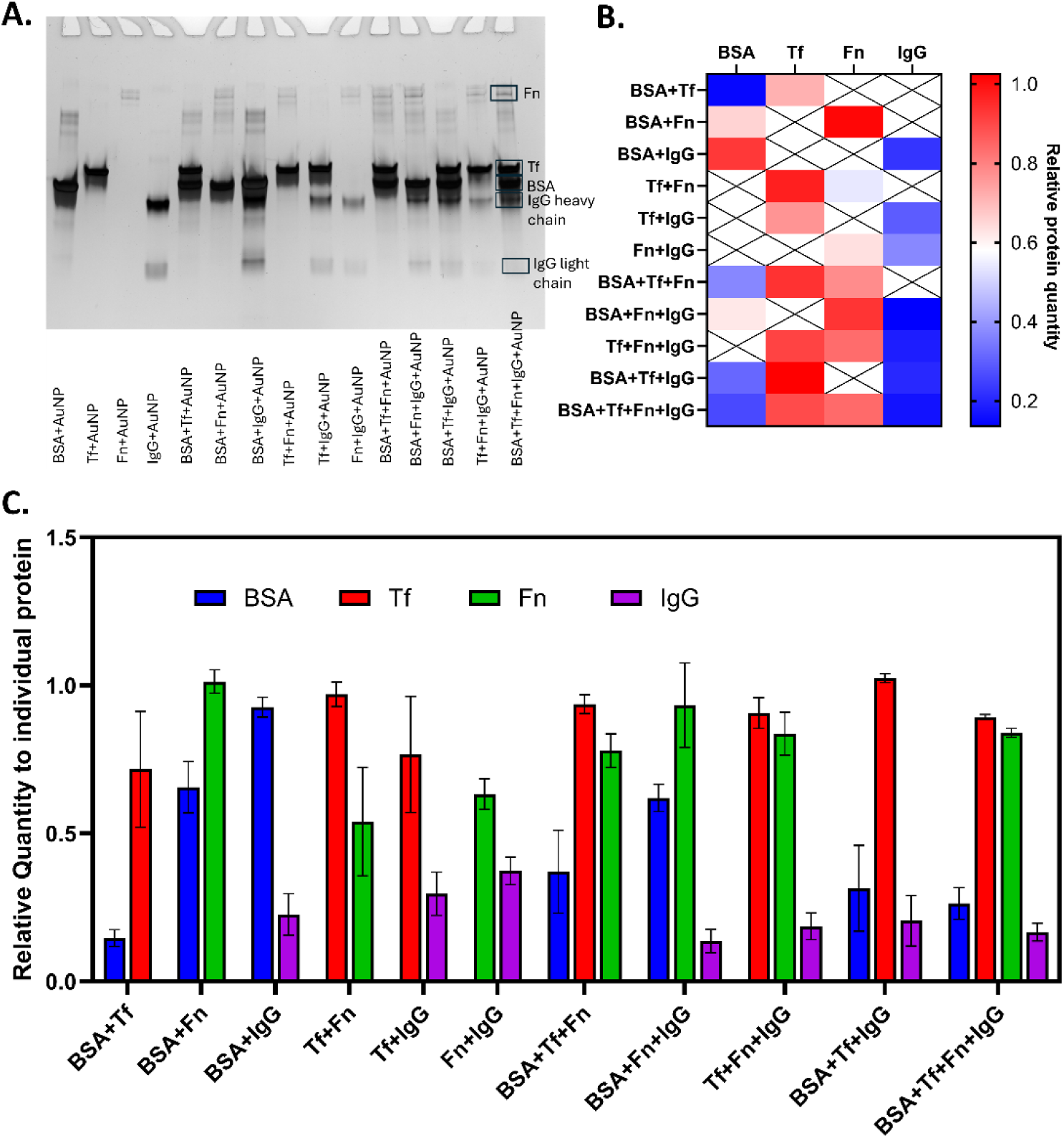
Protein corona composition analysis reveals binding specificity and hierarchical adsorption patterns. (A) SDS-PAGE gel analysis of protein coronas formed on AuNPs with various combinations of BSA, transferrin (Tf), fibronectin (Fn), and immunoglobulin G (IgG). (B) Heat map visualization of relative protein abundance in different corona compositions, with color intensity representing the protein quantity relative to the quantity when incubated alone with nanoparticles. Red indicates higher abundance, blue indicates lower abundance. Proteins that were left out of the original mixture are marked with an X. (C) Quantitative densitometry analysis showing the relative quantity of each protein (normalized to its individual binding) across different protein combinations.

Quantitative densitometry analysis (**Fig. 3C**) further confirmed that Tf and Fn consistently showed higher binding across various protein mixtures. While BSA showed significant binding when paired with IgG alone (BSA+IgG), its adsorption was substantially reduced in the presence of Tf or Fn. Similarly, IgG consistently exhibited the lowest relative adsorption across all mixtures, suggesting it has the lowest binding affinity among the four proteins. These findings demonstrate that protein corona formation follows a hierarchical specificity pattern: Tf ≈ Fn > BSA > IgG. This specificity hierarchy explains our earlier competition observations. Proteins with higher binding affinity (Tf and Fn) effectively out-compete those with lower affinity (BSA and IgG) from the nanoparticle surface. The preferential adsorption of Tf and Fn is particularly significant as these proteins are known to interact with specific cellular receptors that are overexpressed in certain cancer types.

### Structural Integrity of Engineered Protein Coronas: Insights from Circular Dichroism Analysis

To assess whether our engineered corona approach maintains the functional integrity of the constituent proteins, we conducted a detailed circular dichroism (CD) spectroscopy analysis. CD spectroscopy provides valuable information about protein secondary structure, allowing us to determine whether protein-protein interactions or nanoparticle binding induce conformational changes that might alter biological activity. Unlike NMR, CD is unable to monitor individual components in complex mixtures; instead, we examined whether the total CD signal of mixtures was altered in the presence of AuNPs. Then, building from simple mixtures to more complex ones, we could infer the significance of structural perturbations.

Our CD analysis of binary protein mixtures revealed minimal structural perturbations upon mixing. For example, the observed CD spectra of BSA+Tf mixtures closely matched the mathematical sum of their individual spectra (**Fig. 4A**, red vs. blue curves), indicating that these proteins do not substantially influence each other’s conformation in solution. Similarly, the BSA+IgG mixture (**Fig. 4B**) showed preservation of secondary structure elements upon mixing, with no significant deviations between observed and calculated spectra. When these protein mixtures were incubated with AuNPs, we observed subtle but detectable changes in their CD profiles. These changes were quantified by measuring CD signal intensities at 222 nm (**Fig. 4C**) and 208 nm (**Fig. 4D**), wavelengths that are particularly sensitive to α-helical content. Across various protein combinations, the presence of AuNPs consistently induced modest alterations in secondary structure, likely reflecting conformational adaptations as proteins interact with the curved nanoparticle surface.^21^

**Figure 4.**
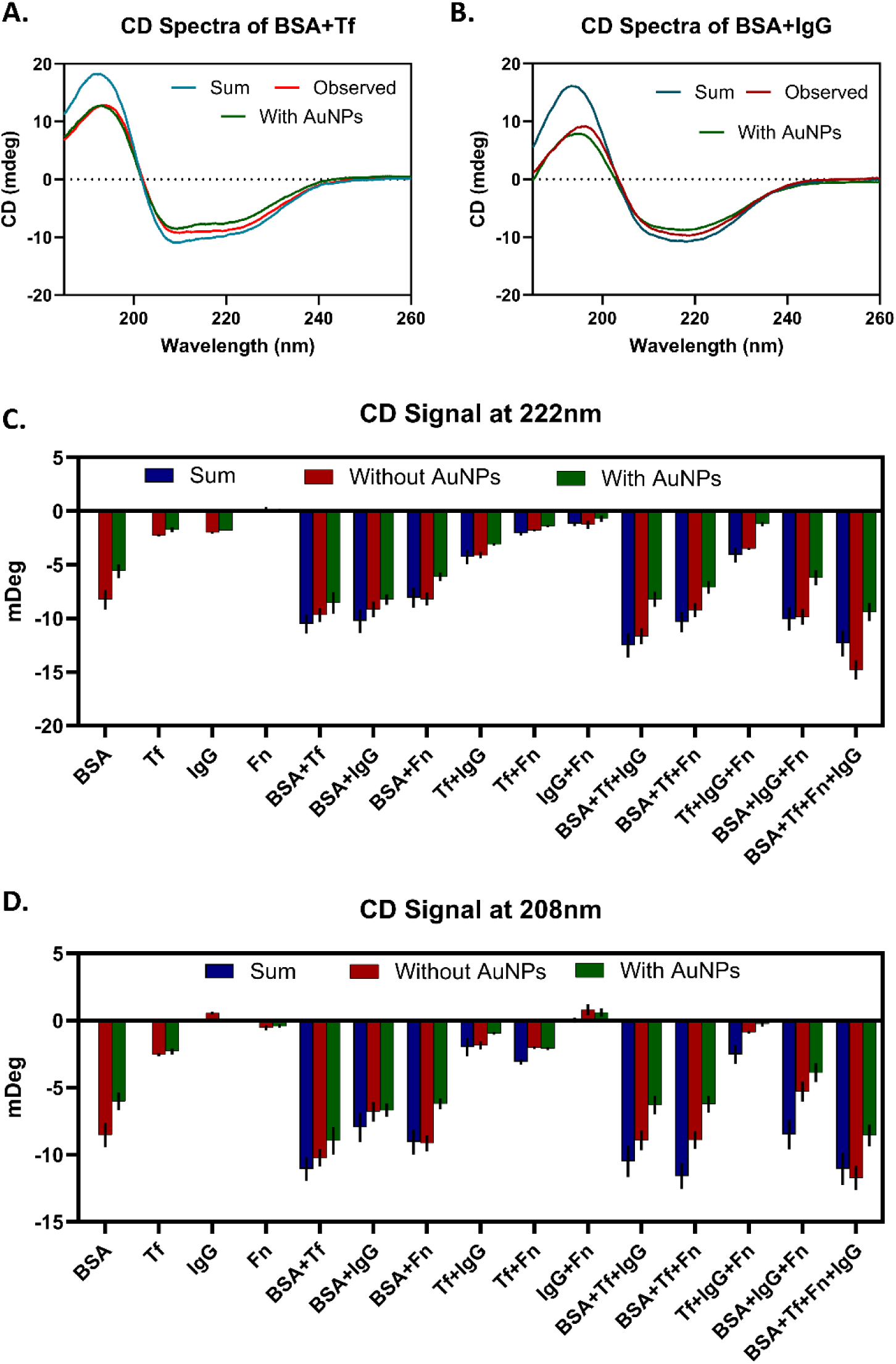
Circular dichroism analysis confirms structural integrity of proteins in engineered coronas. (A) CD spectra of BSA+Tf mixtures showing the mathematical sum of individual protein spectra (blue), the experimentally observed mixture (red), and the mixture with AuNPs (green). (B) Similar CD spectral analysis for BSA+IgG mixtures. (C) Comparative CD signal intensities at 222 nm (α-helical indicator) for various protein combinations, comparing the mathematical sum (blue), observed mixture without AuNPs (red), and with AuNPs (green). (D) Corresponding CD signal analysis at 208 nm across all protein combinations.

Notably, these structural changes were relatively minimal compared to complete protein denaturation, suggesting that proteins in our engineered coronas retain their overall fold and likely their biological function. While we observed structural adaptations in BSA when studied individually with AuNPs, consistent with previous studies showing albumin’s conformational flexibility upon nanoparticle adsorption^61–63^, our data reveal that structural changes occur across various protein combinations, including mixtures without BSA (such as Tf+Fn+IgG). This indicates that surface-induced conformational adaptations are a general phenomenon rather than specific to albumin-containing mixtures.

Our comparative analysis of single, binary, ternary, and quaternary protein combinations revealed that structural changes occur across all levels of corona complexity. Contrary to what might be expected, increasing mixture complexity did not consistently decrease the magnitude of AuNP-induced structural changes. Instead, the CD data suggests that each protein combination interacts uniquely with the nanoparticle surface, resulting in specific conformational adaptations regardless of mixture complexity. For example, while BSA alone experiences a 25% loss in signal when mixed with AuNPs, adding IgG results in a mixture with a stable CD signal in the presence of AuNPs. These CD results are particularly significant for our corona engineering approach, as they demonstrate that while some structural adaptations occur upon nanoparticle binding, our designed protein combinations largely maintain their core structural integrity. This preservation of near-native protein structures suggests that key functional domains, such as the transferrin receptor-binding region or fibronectin’s cell-recognition sequences, may remain sufficiently intact in the engineered corona to maintain their biological activity, which is essential for the targeted interactions we aim to achieve.

Furthermore, the minimal structural changes observed in our complex four-protein corona (BSA+Tf+Fn+IgG) indicate that this biomimetic system closely resembles the behavior of proteins in natural coronas while offering the advantage of a defined composition. This structural robustness provides a solid foundation for rational corona engineering to control nanoparticle behavior in biological environments.

### Engineered Protein Coronas Selectively Enhance Cancer Cell Targeting While Reducing Macrophage Uptake

After establishing the composition and structural integrity of our engineered protein coronas, we investigated their functional impact on cellular interactions. To evaluate their targeting selectivity, we assessed the uptake of various corona-coated AuNPs by MDA-MB-231 triple-negative breast cancer cells and RAW 264.7 macrophages, key cell types that determine therapeutic efficacy and systemic clearance, respectively.

Prior to uptake studies, we assessed the cytotoxicity of all nanoparticle formulations in both cell types at the concentrations used for uptake experiments (Supporting Information, **Fig. S9**). While most formulations showed minimal cytotoxicity with cell viability above 85% after 24-hour exposure at 100 μg mL⁻¹, AuNPs+Fn exhibited moderate cytotoxicity with approximately 70% cell viability in both cell types. This cytotoxic effect may be attributed to fibronectin’s role in cell adhesion and signaling pathways, where high local concentrations on the nanoparticle surface could potentially disrupt normal cellular processes or trigger apoptotic responses.

Bare AuNPs demonstrated modest uptake by both cell types, with approximately 4.7% uptake by macrophages and 2.4% by cancer cells (**Fig. 5A**). When functionalized with single proteins, dramatically different cellular responses emerged. AuNPs coated with IgG showed high macrophage uptake (~26%) with relatively low cancer cell uptake (~13%), consistent with macrophage recognition of the Fc region.^64^ In contrast, AuNPs coated with BSA, Tf, or Fn exhibited a similar targeting profile in cancer cells and macrophages.

**Figure 5.**
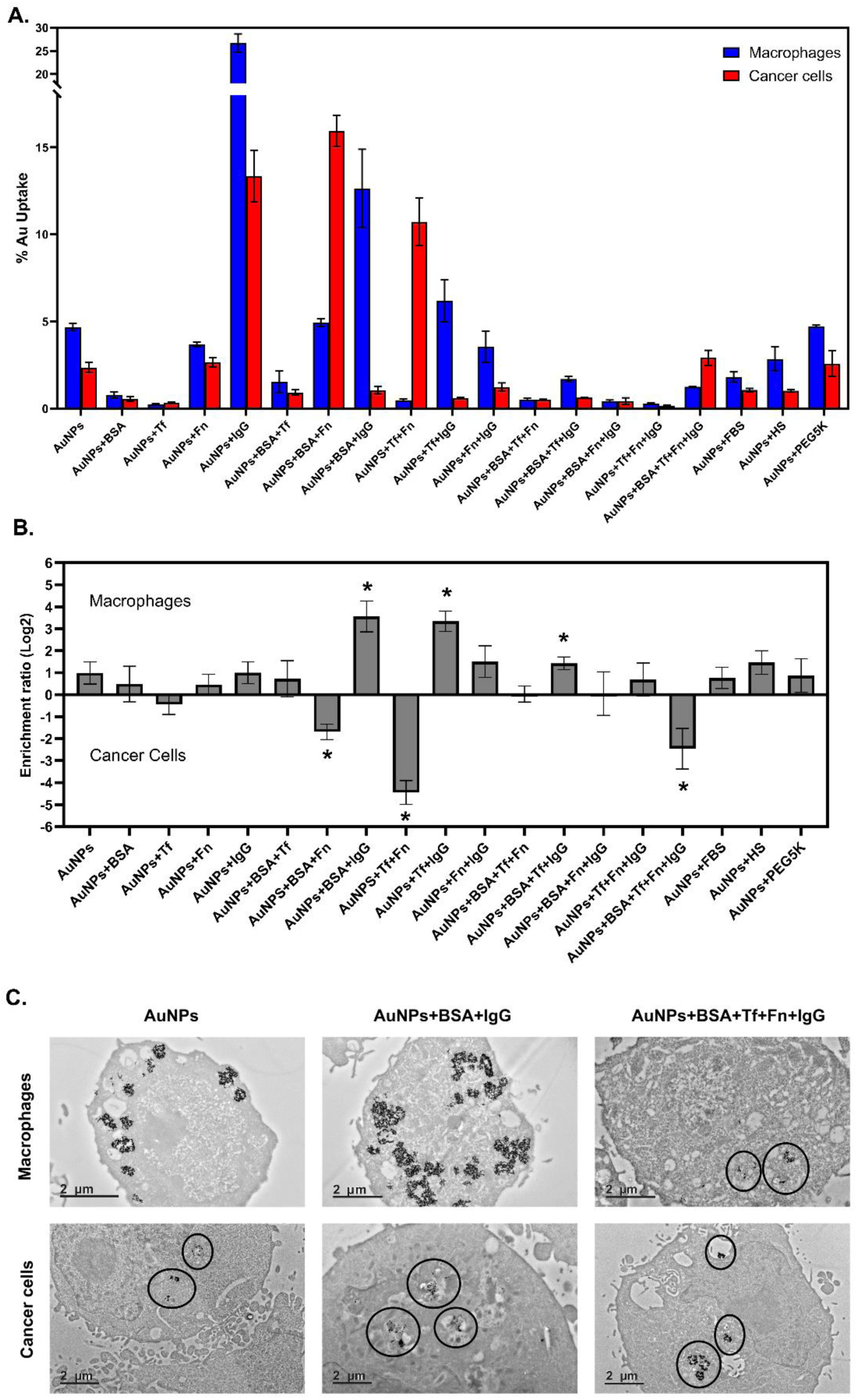
Engineered protein coronas modulate cellular uptake patterns to enhance cancer targeting selectivity. (A) Quantitative uptake analysis of AuNPs with various protein coatings by RAW 264.7 macrophages (blue) and MDA-MB-231 triple-negative breast cancer cells (red), showing differential cellular interactions dependent on corona composition. (B) Cellular enrichment ratios (log₂ scale) for each formulation, with positive values indicating macrophage preference and negative values showing cancer cell selectivity. Asterisks denote statistically significant targeting preferences (p < 0.05). (C) Transmission electron microscopy images of macrophages (top row) and cancer cells (bottom row) after incubation with bare AuNPs, BSA+IgG-coated AuNPs, and AuNPs with the engineered BSA+Tf+Fn+IgG corona. Black circles highlight nanoparticle locations within cells. Scale bars: 2 μm.

The most intriguing findings emerged from analyzing the binary and more complex protein combinations. The BSA+Tf binary mixture showed minimal uptake by both cell types (~1-2%), suggesting a “stealth” property that could potentially extend circulation time. The BSA+Fn formulation demonstrated strong cancer cell selectivity, with approximately 16% cancer cell uptake compared to only 5% macrophage uptake, representing one of the most favorable targeting profiles among all tested coronas. This selectivity could be due to cancer cell overexpression of integrin receptors that recognize fibronectin.^65^ When examining more complex protein combinations, we observed that the four-component formulation (BSA+Tf+Fn+IgG) achieved excellent targeting selectivity with moderate overall uptake (3% cancer versus 1% macrophage), significantly outperforming conventional approaches like PEGylation (~4.7% macrophage, ~2.5% cancer) or FBS coating (~2% macrophage, ~1% cancer).

To better quantify and visualize these targeting preferences, we calculated cellular enrichment ratios (log₂ scale) for each formulation (**Fig. 5B**). This transformation highlights the relative selectivity between the two cell types, with positive values indicating macrophage preference and negative values indicating cancer cell targeting. Six formulations showed statistically significant targeting profiles: AuNPs+BSA+IgG, AuNPs+Tf+IgG and AuNPs+BSA+Tf+IgG strongly favored macrophages, while AuNPs+BSA+Fn, AuNPs+Tf+Fn, and AuNPs+BSA+Tf+Fn+IgG exhibited pronounced cancer cell selectivity.

Transmission electron microscopy provided visual confirmation of these cellular interactions (**Fig. 5C**). Bare AuNPs were observed within both cell types, with evident particle clustering. The BSA+IgG combination showed enhanced macrophage uptake with substantial intracellular accumulation, consistent with the quantitative uptake data. Most significantly, the BSA+Tf+Fn+IgG corona formulation demonstrated selective cancer cell uptake (highlighted by black circles) with minimal presence in macrophages, validating our quantitative findings and confirming the cancer-targeting capability of this engineered corona. Similar targeting selectivity was observed with Tf+Fn-coated AuNPs, while PEGylated AuNPs showed reduced cellular interactions overall (Supporting Information, **Fig. S10**).

These results demonstrate that protein corona composition directly influences cellular interactions in ways that cannot be predicted from individual protein behaviors alone. The competitive protein binding dynamics we observed in our earlier experiments translate into functional consequences at the cellular level. For example, despite IgG’s strong macrophage-activating properties when used alone, its effect is substantially modulated in complex coronas where Tf and Fn demonstrate preferential binding.

These findings align with recent studies on nanoparticle surface composition and immune activation. Building on Park *et al*.’s work on “cloaking” nanoparticles with functional protein coatings,^66^ our approach extends beyond simple immune evasion to create a corona with multiple complementary functions. While Park *et al*. demonstrated that specific proteins like complement factor H could reduce complement activation and macrophage uptake, our engineered corona incorporates additional targeting capabilities toward cancer cells, representing an advancement in rational protein corona design. Our findings also align with Mirshafiee *et al*.’s observation that protein orientation and accessibility within the corona are crucial for maintaining targeting capabilities.^33^ However, where they found that opsonin-rich coronas failed to enhance macrophage uptake due to steric hindrance, our multi-protein engineering approach successfully maintained targeting specificity, demonstrating advantages over single-protein pre-coating strategies.

Our approach offers significant advantages over conventional strategies. While PEGylation is widely used to reduce non-specific uptake, it lacks intrinsic targeting capabilities and often suffers from the “PEG dilemma,” where reduced macrophage recognition comes at the cost of reduced overall cellular interactions.^67^ PEGylation is also increasingly associated with undesirable immune responses.^68, 69^ In contrast, our engineered corona approach can simultaneously reduce macrophage uptake while enhancing cancer cell targeting through receptor-specific interactions.

### Proteomics Reveals Distinctive Secondary Corona Formation on Engineered Nanoparticles

After establishing the targeting capabilities of our engineered coronas in vitro, we sought to understand why and how these pre-formed protein layers interact with the complex protein milieu encountered upon systemic administration. We selected representative formulations showing distinct cellular targeting behaviors, cancer-targeting (BSA+Fn, Tf+Fn, BSA+Tf+Fn+IgG) and macrophage-targeting (BSA+IgG, Tf+IgG, BSA+Tf+IgG) formulations, along with simpler non-engineered controls (bare AuNPs, PEGylated AuNPs) for comprehensive proteomics analysis. We employed quantitative proteomics to characterize the “secondary corona” that forms when our engineered nanoparticles interact with human serum, providing critical insights into their biological identity and potential in vivo behavior. By characterizing the secondary corona, we aimed to identify additional proteins that may be recruited and that may influence uptake.

All nanoparticle formulations were incubated with human serum under physiological conditions, followed by isolation and characterization of the resulting protein corona. SDS-PAGE analysis (**Fig. 6A**) revealed distinctive protein pattern differences across formulations, with visible variations in band intensity and distribution, confirming that the pre-formed engineered coronas lead to formulation-specific serum protein adsorption. DLS measurements (**Fig. 6B**) provided quantitative evidence of differential protein adsorption behavior across formulations. While most formulations exhibited increased hydrodynamic diameters ranging between approximately 200-280 nm after serum exposure, several specific formulations showed significantly reduced dimensions compared to bare AuNPs (~230 nm), indicating their distinctive interactions with serum proteins. AuNPs+PEG5K showed the most dramatic reduction in hydrodynamic diameter to approximately 50 nm, consistent with its known “stealth” properties in biological systems. Similarly, AuNPs+BSA+Tf demonstrated a significant reduction in diameter to approximately 100 nm, suggesting this specific protein combination creates a corona that effectively limits secondary protein adsorption.

**Figure 6.**
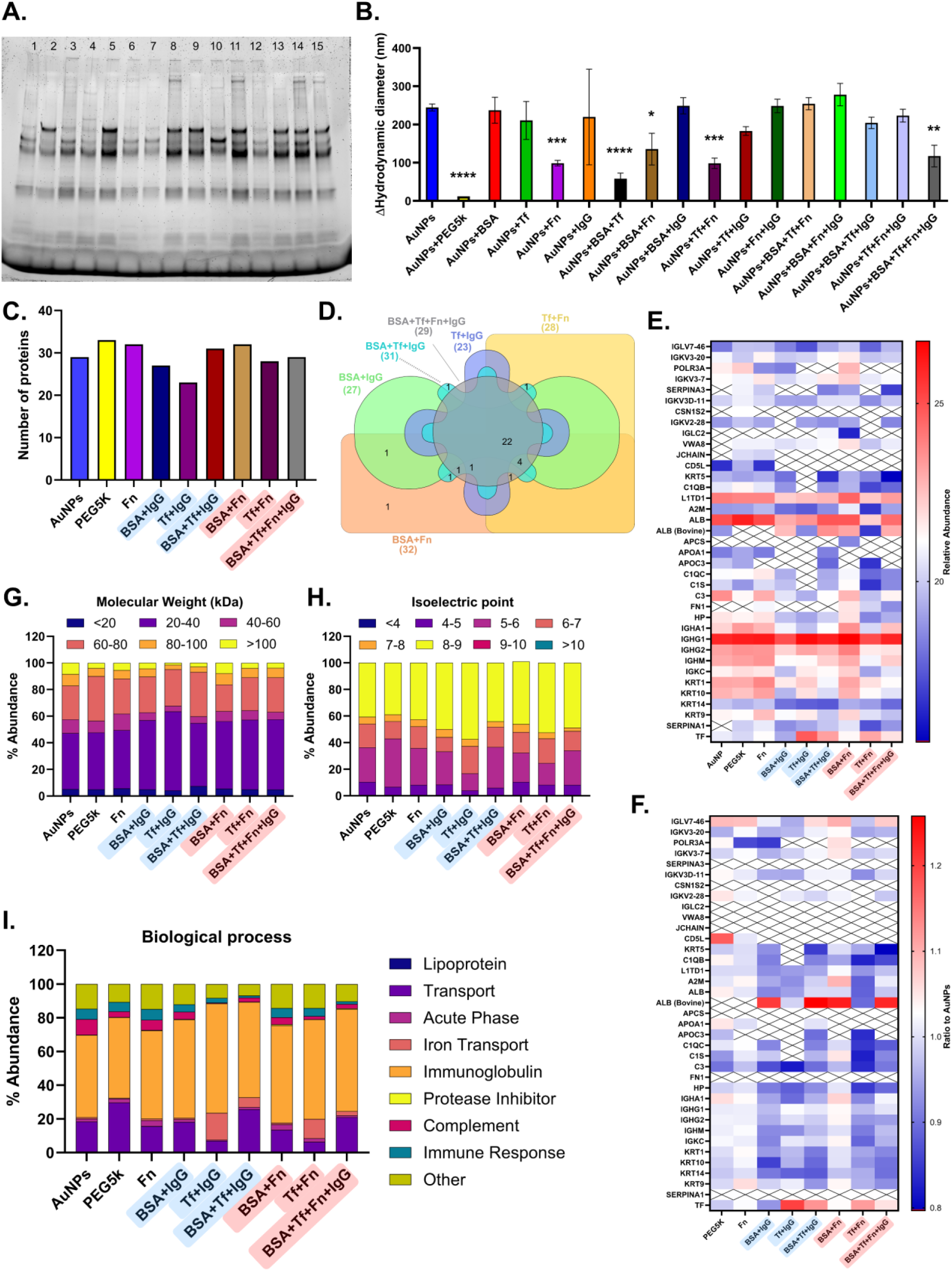
Proteomics analysis reveals distinct protein corona signatures on engineered nanoparticles after serum exposure. (A) SDS-PAGE analysis of protein coronas formed on different AuNP formulations after incubation with human serum, showing formulation-dependent band patterns. Lane assignments: 1) AuNPs+BSA, 2) AuNPs+Tf, 3) AuNPs+Fn, 4) AuNPs+IgG, 5) AuNPs+BSA+Tf, 6) AuNPs+BSA+Fn, 7) AuNPs+BSA+IgG, 8) AuNPs+Tf+Fn, 9) AuNPs+Tf+IgG, 10) AuNPs+Fn+IgG, 11) AuNPs+BSA+Tf+Fn, 12) AuNPs+BSA+Fn+IgG, 13) AuNPs+BSA+Tf+IgG, 14) AuNPs+Tf+Fn+IgG, 15) AuNPs+BSA+Tf+Fn+IgG. (B) The increase in the hydrodynamic diameter of various AuNP formulations after exposure to human serum. Statistical significance was assessed using one-way ANOVA followed by Dunnett’s multiple comparisons test with AuNPs as the reference group. Asterisks indicate significant differences compared to AuNPs control: *p < 0.05, **p < 0.01, ***p < 0.001, ****p < 0.0001. (C) Quantitative analysis of the total number of unique proteins identified by LC-MS/MS across different nanoparticle formulations. (D) Overlap analysis (Venn diagram) showing common proteins between macrophage-targeting and cancer cell-targeting formulations. (E) Relative abundance of serum proteins identified by LC-MS/MS across different nanoparticle formulations, with color intensity representing protein abundance (red: higher, blue: lower). (F) Heatmap visualization of protein abundance ratios normalized to bare AuNPs, highlighting distinctive protein recruitment signatures across formulations. Color intensity corresponds to the normalized abundance ratio (red: higher than AuNPs, blue: lower than AuNPs, x: protein not detected). (G) Classification of corona proteins by molecular weight (kDa), (H) isoelectric point (pI), and (I) biological process (Gene Ontology analysis).

Other formulations showing statistically significant reductions in hydrodynamic diameter compared to bare AuNPs included AuNPs+Fn (~140 nm), AuNPs+Tf+Fn (~100 nm), AuNPs+BSA+Fn (~200 nm), and AuNPs+BSA+Tf+Fn+IgG (~125 nm). These findings extend our earlier competition studies by revealing how our engineered coronas not only exhibit specific protein binding hierarchies but also influence the extent and composition of secondary corona formation when exposed to complex protein mixtures. The moderate reduction observed with our BSA+Tf+Fn+IgG formulation indicates a unique mode of interaction with serum proteins, neither completely blocking protein adsorption (as with PEGylation) nor permitting unrestricted protein binding (as with simple protein coronas), but rather selectively modulating the secondary corona formation.

The diversity of protein recruitment across formulations was examined by quantifying the total number of unique proteins identified for each nanoparticle system (**Fig. 6C**). Most formulations recruited between 23-33 detectable proteins, with relatively modest variation across different corona compositions. Notably, BSA+Tf+Fn+IgG formulation bound 29 proteins, similar to bare AuNPs, while PEGylated AuNPs showed the highest protein count (33). This suggests that corona engineering primarily modulates protein identity and abundance rather than simply reducing protein diversity, challenging the assumption that surface modifications necessarily decrease protein interactions. The result for PEGylated AuNPs also establishes that protein recruitment in proteomics does not necessarily correspond to a significant increase in hydrodynamic diameter.

To understand the molecular basis of differential cellular targeting, we analyzed protein overlap between formulations showing distinct biological behaviors (**Fig. 6D**). Remarkably, cancer-targeting formulations (highlighted in red) and macrophage-targeting formulations (highlighted in blue) shared 22 common proteins out of their total proteomes (29-32 proteins each). This substantial overlap indicates that targeting specificity does not arise from recruiting fundamentally different sets of proteins, but rather from how these shared proteins are organized, oriented, and presented on the nanoparticle surface in the context of the pre-formed corona.

Quantitative analysis of protein abundance (**Fig. 6E**, Supporting Information **Table S1**) revealed both consistencies and important variations across formulations. Key serum proteins, including albumin (ALB), immunoglobulins (IGHG1, IGHG2), and complement components, showed high abundance across most formulations. However, targeting proteins showed formulation-specific variations: transferrin (TF) demonstrated higher levels in BSA+Tf+Fn+IgG compared to bare AuNPs, confirming retention of pre-bound targeting proteins.

To better visualize formulation-specific protein affinities, we calculated abundance ratios relative to bare AuNPs for each protein (**Fig. 6F**, Supporting Information **Table S2**). This normalization revealed distinctive protein recruitment signatures across formulations. Transferrin showed enhanced retention in Tf-containing formulations (ratios 1.05-1.20), while complement protein C3 demonstrated reduced binding to several engineered coronas (ratios 0.84-0.97), with BSA+Tf+Fn+IgG showing a ratio of 0.90. This reduced complement association may contribute to improved circulation properties. Apolipoproteins exhibited selective recruitment patterns, with APOA1 showing enhanced binding to PEG5K (ratio 1.05) compared to other formulations, suggesting that different surface chemistries can selectively recruit specific lipoproteins. The heatmap visualization illustrates how different formulations create unique protein signatures that correlate with their observed biological behaviors.

To understand the physicochemical basis underlying these protein recruitment patterns, we analyzed the molecular characteristics of corona proteins across formulations. Molecular weight distribution analysis (**Fig. 6G**) revealed that all formulations recruited proteins spanning the full range from small (<20 kDa) to large (>100 kDa) proteins, with the 20-40 and 60-80 kDa range representing the dominant fraction across most formulations. This size range includes many key serum proteins such as albumin (66 kDa) and transferrin (80 kDa). Notably, cancer cell-targeting formulations showed slightly enhanced recruitment of proteins in the 80-100 kDa range, which encompasses important targeting molecules like transferrin and fibronectin fragments, while maintaining moderate levels of larger immunoglobulins (>100 kDa).

Isoelectric point analysis (**Fig. 6H**) revealed distinct pH-dependent recruitment patterns between targeting modalities. Cancer-targeting formulations showed enhanced recruitment of acidic proteins (pI 4-5), including many important transport and cell adhesion proteins such as transferrin (pI ~5.4) and certain fibronectin fragments. This acidic protein preference may facilitate interactions with cancer cells, which often exhibit altered surface charge distributions and modified extracellular pH environments compared to normal tissues. In contrast, macrophage-targeting formulations demonstrated broader pI distribution extending into more basic regions (pI 8-9), potentially reflecting recruitment of complement components, antimicrobial proteins, and certain immunoglobulins that facilitate immune recognition. The more neutral pI profile (6-7) was moderately represented across all formulations, corresponding to abundant serum proteins like albumin.

The functional classification analysis (**Fig. 6I**) provided the most direct link between protein corona composition and observed biological behaviors. Across all formulations, immunoglobulins represented the largest functional category, followed by protease inhibitors and transport proteins. However, critical differences emerged in the relative proportions of functional classes. Cancer cell-targeting formulations, particularly BSA+Tf+Fn+IgG, showed enhanced recruitment of transport proteins and reduced proportion of immune response and complement proteins compared to macrophage-targeting formulations like BSA+IgG.

PEGylated AuNPs displayed a unique functional profile with minimal complement proteins and reduced immune response components, consistent with their established stealth properties. Acute phase response proteins were present across most formulations but in varying proportions, with cancer-targeting formulations showing more balanced representation compared to the immune-skewed profiles of macrophage-targeting systems.

This functional analysis provides a molecular explanation for our in vitro cellular targeting results: formulations enriched in transport and cell adhesion proteins achieved enhanced cancer cell recognition through receptor-mediated interactions, while those with higher immune response protein content triggered increased macrophage uptake through opsonization pathways. The distinct protein corona signature of our BSA+Tf+Fn+IgG formulation, combining sufficient transport protein recruitment with reduced immune activation signals, and maintained levels of targeting proteins (Tf, Fn) creates an ideal balance for selective cancer targeting while evading macrophage recognition. Notably, correlation analysis reveals that cancer targeting is not simply determined by individual protein abundance levels, but rather by the synergistic interactions between multiple targeting proteins within the engineered corona (Supporting Information, **Fig. S11**).

### Engineered Protein Coronas Exhibit Distinctive Pharmacokinetic Profiles and Enhanced Tumor Accumulation In Vivo

To evaluate the in vivo performance of our engineered corona approach, we investigated the pharmacokinetics and biodistribution of various nanoparticle formulations in 4T1 breast cancer tumor-bearing mice. We hypothesized that the enhanced cancer targeting and reduced macrophage uptake observed in vitro would translate to improved circulation time and tumor accumulation in vivo.

We first assessed the blood circulation profiles of different nanoparticle formulations following intravenous administration in BALB/c mice bearing subcutaneous 4T1 breast cancer tumors. The gold content in blood samples was quantified using inductively coupled plasma mass spectrometry (ICP-MS) over a 24-hour period, according to the experimental design outlined in **Fig. 7A**. As shown in **Fig. 7B**, the formulations demonstrated markedly different pharmacokinetic behaviors. Bare AuNPs were rapidly cleared from circulation, with gold concentration dropping below 20 ppm within 6 hours. Single-protein coronas showed variable circulation behavior, with BSA+IgG-coated nanoparticles exhibiting the most rapid clearance, while BSA+Fn-coated nanoparticles demonstrated significantly extended circulation.

**Figure 7.**
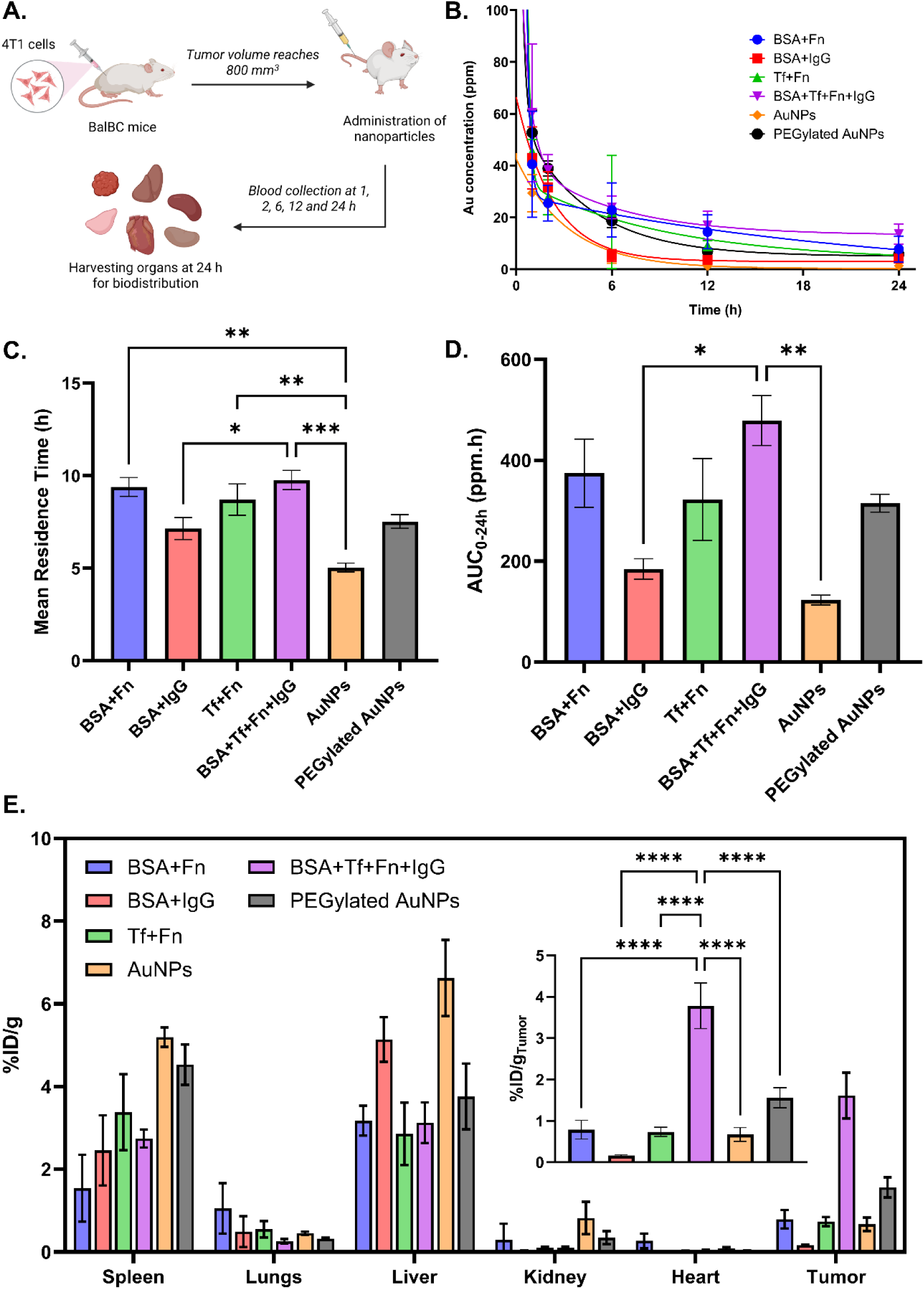
In vivo pharmacokinetics and biodistribution demonstrate enhanced tumor targeting with engineered protein coronas. (A) Schematic illustration of the experimental design using 4T1 tumor-bearing BALB/c mice, including administration of nanoparticles, blood collection time points, and organ harvesting for biodistribution analysis. (B) Blood circulation profiles of various AuNP formulations measured over 24 hours. (C) Calculated mean residence time for each formulation in circulation. (D) Area under the curve (AUC_0-24h_) analysis quantifying total nanoparticle exposure over 24 hours. (E) Biodistribution of nanoparticles in major organs and tumor tissue at 24h post-administration, expressed as gold concentration per gram of tissue (ppm/g), with the inset highlighting tumor accumulation. Error bars represent SEM. Asterisks indicate statistical significance determined using a one-way ANOVA and Tukey’s honestly significant difference post-hoc analysis. (*p < 0.05, **p < 0.01, ****p < 0.0001).

Initial attempts to characterize elimination kinetics using conventional two-compartment models proved problematic, yielding inconsistent parameter estimates and poor goodness-of-fit statistics (R² = 0.59-0.96). This instability likely reflects the inherently complex clearance mechanisms of engineered protein corona systems, which may undergo dynamic protein exchange, sequential recognition events, and multi-organ distribution processes that violate standard pharmacokinetic assumptions. Given these limitations, we employed mean residence time (MRT) as our primary circulation metric, providing a robust, model-independent assessment of systemic persistence.

Non-compartmental analysis revealed a complex relationship between corona complexity and circulation kinetics (**Fig. 7C**). The BSA+Tf+Fn+IgG formulation achieved the longest MRT (9.77 ± 0.52 h), representing a 94% improvement over bare AuNPs (5.04 ± 0.24 h) and a 30% enhancement compared to PEGylated controls (7.53 ± 0.37 h). BSA+Fn (9.39 ± 0.51 h) and Tf+Fn (8.71 ± 0.85 h) both exceeded PEGylated benchmarks by 25% and 16%, respectively. Notably, BSA+IgG (7.15 ± 0.59 h) provided minimal circulation enhancement, suggesting that protein identity, not merely presence, determines stealth efficacy.

Area under the curve (AUC_0-24h_) analysis (**Fig. 7D**) provided insights into the overall systemic exposure. The BSA+Tf+Fn+IgG formulation demonstrated the highest AUC (~500 ppm·h), significantly greater than all other formulations, including BSA+Fn, Tf+Fn, and PEGylated AuNPs. Bare AuNPs showed the lowest AUC (approximately 130 ppm·h), consistent with their rapid clearance. These findings indicate that the BSA+Tf+Fn+IgG formulation exhibits a complex pharmacokinetic profile with longest MRT and maintaining sufficient circulation to achieve superior overall exposure.

Biodistribution analysis at 24 hours post-injection revealed distinct accumulation patterns across formulations (**Fig. 7E**). All formulations showed substantial liver accumulation, consistent with the liver being a primary clearance organ for nanoparticles. AuNPs and BSA+IgG-coated nanoparticles showed the highest liver accumulation, while BSA+Tf+Fn+IgG and BSA+Fn demonstrated moderately lower liver uptake (approximately 3% ID/g). Spleen accumulation was highest for bare AuNPs (approximately 5% ID/g) and lowest for BSA+Fn (approximately 1.5% ID/g), reflecting differences in recognition by the reticuloendothelial system. Low accumulation was observed in lungs, kidneys, and heart across all formulations, as expected for nanoparticles of this size range. The most striking finding emerged when examining tumor accumulation (inset, **Fig. 7E**). The BSA+Tf+Fn+IgG formulation achieved remarkably high tumor accumulation of approximately 4% ID/g (13 ppm/g), significantly higher than all other formulations (Supporting Information, **Fig. S11**). This represents a 6.5-fold improvement compared to bare AuNPs (~0.8% ID/g), a 26-fold improvement over BSA+IgG (~0.2% ID/g), a 4.3-fold improvement over BSA+Fn (~1% ID/g), a 5.2-fold improvement over Tf+Fn (~1% ID/g), and a 2.6-fold improvement over PEGylated AuNPs (~1.5% ID/g). These differences were statistically significant (p < 0.0001), underscoring the superior tumor targeting capability of our engineered four-protein corona.

The enhanced tumor accumulation of the BSA+Tf+Fn+IgG formulation is particularly noteworthy given that it achieved both extended circulation and the highest tumor targeting efficiency. This dual advantage suggests that this formulation succeeds through a combination of mechanisms: prolonged systemic exposure that maximizes opportunities for tumor encounter, coupled with active targeting mechanisms that enhance tumor cell recognition and retention. The presence of both Tf and Fn in this corona likely contributes to enhanced tumor cell recognition and retention, as these proteins can interact with transferrin receptors and integrins that are overexpressed on cancer cells. Additionally, the specific arrangement of these proteins in the engineered corona appears to provide an optimal balance between immune evasion (reducing liver and spleen accumulation) and active targeting (enhancing tumor uptake). The superior performance of our BSA+Tf+Fn+IgG formulation compared to PEGylated AuNPs is particularly significant from a translational perspective. While PEGylation achieved its intended stealth function with reduced liver and spleen uptake compared to bare AuNPs, our engineered corona approach demonstrated comparable organ distribution profiles (similar liver and spleen accumulation) while achieving 2.6-fold higher tumor accumulation. This suggests that rational protein corona engineering may offer advantages over traditional surface modification strategies by harnessing the natural biological interactions of proteins rather than attempting to shield the nanoparticle surface using PEG.

It is important to note that the tumor accumulation achieved by our BSA+Tf+Fn+IgG formulation (approximately 13 ppm/g, equivalent to about 4% ID/g) significantly exceeds the median delivery efficiency of 0.7% ID/g reported for nanomedicines in a comprehensive analysis by Wilhelm et al.^3^ This improvement in delivery efficiency addresses one of the fundamental challenges limiting clinical translation of nanomedicines for cancer therapy.

The correlation between our in vitro findings and in vivo performance supports the validity of our corona engineering approach. The BSA+Tf+Fn+IgG formulation, which demonstrated enhanced cancer cell targeting and reduced macrophage uptake in vitro, achieved the most efficient tumor targeting in vivo. This consistency across experimental platforms suggests that rational corona engineering can effectively influence nanoparticle-biological interactions at multiple scales, from molecular binding events to whole-organism distribution patterns.

The in vivo performance of our BSA+Tf+Fn+IgG formulation results from a coordinated mechanism that integrates hierarchical protein competition, structural preservation, and functional protein recruitment. Our data reveal that high-affinity proteins (Tf, Fn) preferentially dominate the corona composition through competitive interactions, while CD analysis confirms these proteins retain their structural integrity and targeting functionality upon surface binding. This preserved biological activity enables the engineered corona to influence secondary protein recruitment when exposed to serum, creating distinctive functional signatures enriched in transport and cell adhesion proteins rather than immune recognition molecules. The resulting protein profile facilitates selective cancer cell interactions through multiple receptor-mediated pathways (transferrin receptors, integrins) while maintaining balanced immune protein levels that reduce macrophage recognition and complement activation.

This integrated mechanism explains why our BSA+Tf+Fn+IgG formulation achieves both extended circulation (9.77 h MRT) and exceptional tumor accumulation (6.5-fold improvement over bare nanoparticles). Rather than relying on passive stealth properties or single targeting mechanisms, the engineered corona creates a “smart” biological interface that simultaneously evades immune clearance and actively targets cancer cells through coordinated protein interactions. The correlation between our molecular-level protein recruitment data and in vivo biodistribution validates this approach as a rational strategy for enhancing nanoparticle targeting by harnessing, rather than avoiding, the natural protein corona formation process to achieve predictable therapeutic delivery.

## Conclusion

This work demonstrates that rational protein corona engineering can address fundamental limitations in nanomedicine delivery through precise control of nanoparticle-biological interactions. By establishing a hierarchical protein binding model (Tf ≈ Fn > BSA > IgG), we created engineered coronas that achieve predictable biological targeting rather than passive stealth behavior. The BSA+Tf+Fn+IgG formulation achieved 4% ID/g tumor accumulation—nearly 6-fold higher than current clinical benchmarks and surpassing PEGylated controls by 2.6-fold.

This enhanced targeting results from coordinated protein interactions that simultaneously reduce immune recognition while promoting cancer cell uptake through multiple receptor pathways. Proteomics analysis reveals that effective targeting relies on recruiting transport and cell adhesion proteins while minimizing immune activation signatures, creating biological interfaces that harness rather than resist natural corona formation. These findings establish protein corona engineering as a new avenue for nanomedicine design. Rather than attempting to shield nanoparticles from biological recognition, our approach exploits the corona as a programmable targeting system.

Moreover, while these results are highly promising, they likely represent only an early benchmark, as further optimization of the formulation could yield substantially greater performance. The mechanistic principles demonstrated here—hierarchical protein competition, preserved targeting functionality, and selective secondary protein recruitment—provide a rational framework for designing next-generation nanomedicines with improved therapeutic efficacy.

## Materials and Methods

### Synthesis of citrate-capped AuNPs

AuNPs with a diameter of 15 nm were synthesized using the citric acid reduction method based on the Turkevich synthesis.^70^ Tetrachloroauric acid (HAuCl_4_, #520918) and sodium citrate dihydrate (#W302600) were obtained from Millipore-Sigma. First, 2 mL of HAuCl_4_ solution (10 mg mL^−1^) was mixed with 198 mL of water and heated to a boil. This mixture was allowed to rest for 10 min, and then 7 mL of sodium citrate solution (10 mg mL^−1^ of the dihydrate) was added. The mixture was stirred at 400 RPM and heated for 25 minutes at power level 48 (power level refers to the percentage of maximum output voltage of 120 V) before stirring was stopped. The sample was allowed to cool until reaching room temperature. Nanoparticles were then concentrated by centrifugation at 9,000 x g for 45 min. AuNPs were then characterized using UV-visible (UV-Vis) spectroscopy and dynamic light scattering (DLS, see Supporting Information Figures S1 and S2). The UV-Vis spectrum and DLS-determined diameter were consistent with previous values for 15 nm AuNPs.^38, 71^ The expected maximum absorbance of the 15 nm AuNPs was determined to be at 520 nm with an extinction coefficient of 3.9×10^8^ M^−1^ cm^−1^.^72^ AuNPs were used within a week after synthesis.

### Preparation of protein mixtures

Fetal bovine serum (FBS) was purchased from Millipore-Sigma (#F2442, lot no. 21G638) and stored at −20 °C. To limit freeze-thaw cycles, FBS was aliquoted into 1 mL tubes before freezing. Bovine serum albumin (BSA, #12657), human plasma fibronectin (Fn, #FC010), human transferrin (Tf, #T8158), and Immunoglobulin G (IgG, #I4506) were obtained from Millipore-Sigma and used without further purification. Individual protein samples were prepared by dissolving the lyophilized powder in 20 mM sodium phosphate buffer (pH 7.4). The mixture containing all proteins was created, where protein concentration ratios matched those found in serum.^73, 74^ For a BSA concentration of 1 mg mL^−1^, 0.1 mg mL^−1^ Tf, 0.05 mg mL^−1^ Fn, and 0.6 mg mL^−1^ IgG were used, for a total protein concentration of approximately 1.8 mg mL^−1^. These specific values represent a dilute total protein concentration compared to normal serum, which typically contains 30-50 mg mL^−1^ BSA. When a more (or less) concentrated mixture was used, all concentrations were scaled up (or down) accordingly. This design ensures that proteins with lower absolute concentrations that successfully compete for binding demonstrate genuine binding preference rather than concentration-driven effects. Different combinations of these proteins were mixed to make a simple mixture, and all the proteins were mixed to form a more complex mixture as specified in the text.

## 13C methylation of BSA and proline

Formaldehyde (^13^C labeled) was purchased from Cambridge Isotope Laboratories (#CLM-806-1), and dimethylamine borane complex was obtained from Thermo Scientific (#AA8914914). 40 µL of 1.0 M formaldehyde was added to 1.0 mL of BSA solution in phosphate buffer (10 mg mL^−1^), and 20 µL of the 1.0 M dimethylamine borane solution was added to the mixture. The mixture was then gently mixed and incubated at 4 °C for 2 hr. After the first 2-hour incubation, an additional 20 µL of the dimethylamine borane solution was added along with 40 µL of 1.0 M formaldehyde. This second addition ensured that the reaction was nearly complete. The mixture was again gently mixed and incubated at 4 °C for another 2 hr. After the second incubation, another 10 µL of dimethylamine borane was added, and the sample was incubated at 4 °C overnight. The methylation was stopped by adding 125 µL of 1.0 M glycine and incubating the mixture at 4 °C for 2 hr. The protein was purified using a Clarion P-10 gel filtration column (Sorbent Technologies). An identical procedure was used to methylate the imine position of a 1 mM proline solution. To avoid issues with batch-to-batch reproducibility, the same methylated proline solution was used as a quantitative standard for all NMR experiments presented in this work.^75, 76^

## 1D proton experiments

1D proton experiments were used to determine the binding capacity of BSA on AuNPs. For these experiments, a 20 µM BSA solution was prepared in 20 mM sodium phosphate buffer, 50 mM NaCl at pH 7.4. These protein samples were mixed with AuNPs at concentrations from 10 to 100 nM. These samples were incubated at room temperature for 2 hours to allow the proteins to bind to AuNPs. 1D proton spectra were recorded on a 600 MHz Bruker Avance III cryoprobe-equipped NMR spectrometer. The recycle delay was set to 1.5 sec, and an acquisition time of 80 ms was used. The spectrum was signal-averaged over 512 scans using the one-one echo experiment for water suppression.^51^ The total experiment time for each sample was 14 mins. All the 1D NMR spectra were processed and baseline corrected using Bruker TOPSPIN software. The AuNP binding capacity (number of BSA bound per AuNP) was determined as described previously.^51, 77^

## 2D ^1^H-^13^C HSQC experiments

HSQC experiments were used to investigate the binding capacity and to determine how serum proteins influence the AuNP adsorption of methylated BSA. The samples for measuring binding capacity were made using the same method described above, but with ^13^C-methylated BSA. A 1 mg mL^−1^ solution of ^13^C-methylated BSA was mixed with different concentrations of unlabeled serum, or mixtures of unlabeled serum proteins, and incubated along with 100 nM AuNPs for 2 hr at room temperature. ^13^C methylated proline was used as an external quantitative standard in a coaxial insert tube (Wilmad #WGS-5BL) along with 50% D_2_O and 20 mM phosphate buffer (pH 7.4). ^1^H-^13^C HSQC spectra were recorded on the same 600 MHz NMR spectrometer with an acquisition time of 170 ms in the indirect dimension (256 complex points) and a total experiment time of 29 min (with 2 scans per increment and a recycle delay of 1 s). The indirect sweep width was 10 ppm. The acquired NMR data were processed using NMRPipe, and the assignment and integration were done using Sparky.^78^ Both dimensions were zero-filled with at least three additional powers of two beyond the number of acquisition points. This ensures the lineshapes are smooth and maximizes the accuracy of relative peak intensities.^48^

### Particle size and zeta potential measurements

All dynamic light scattering (DLS) measurements were performed on an Anton Paar Litesizer 500 instrument using Kalliope software. The samples were prepared as mentioned above, and after equilibrating for 2 h, the solutions were diluted five times and transferred into disposable cuvettes for the DLS measurement. The zeta potential of AuNPs under different conditions was also measured using the Litesizer 500. The samples were prepared and transferred to the Anton Paar Omega cuvette Z (#225288) for measurements.

### SDS-PAGE and densitometry analysis

The protein mixtures were prepared as mentioned above and incubated with 100 nM AuNPs for 2 h. These samples were washed three times to remove any unbound protein. To the protein-coated AuNPs, 50% human serum was added and incubated at room temperature for 1 h. Following serum incubation, samples were washed three times to remove unbound serum proteins. These samples were mixed with 2X Laemmli sample buffer and heated at 95°C for 5 minutes. Samples were loaded on a 4-20% Mini Protean precast TGX gel along with a prestained ladder. Gels were run in 1X SDS-PAGE running buffer at 160 V until the dye front reached the bottom. After electrophoresis, the gels were stained with oriole stain and imaged using a Chemidoc MP imaging system. Digital gel images were analyzed using Image Lab software. Lanes and bands were manually selected. The molecular weights of bands were determined based on the protein ladder. The densitometry analysis quantified the relative intensity of each protein band in a mixture relative to its initial intensity.

### Liquid chromatography-tandem mass spectroscopy (LC-MS/MS) analysis

#### In-gel digestion

In-gel digestion was performed essentially as described by Shevchenko et al. with minor modifications.^79^ Protein bands were excised, destained with 100 mM ammonium bicarbonate in 50% acetonitrile, dehydrated in neat acetonitrile, and dried under vacuum. Gel pieces were reduced with 50 mM dithiothreitol (DTT) at 37 °C for 1–2 h, then alkylated with 55 mM iodoacetamide in 100 mM ammonium bicarbonate for 30–45 min at room temperature in the dark. Alkylation was quenched by addition of DTT to a final concentration of 10 mM. After dehydration, gel pieces were rehydrated in sequencing-grade trypsin solution (10 ng µL^−1^ in 10 mM ammonium bicarbonate, 10% acetonitrile) on ice for 30–90 min, then covered with 100 mM ammonium bicarbonate and digested overnight at 37 °C. Peptides were extracted sequentially with 50% acetonitrile/5% formic acid and 75% acetonitrile/0.1% formic acid, pooled, and dried under vacuum. Pellets were resuspended in LC–MS loading buffer (98:2 H₂O:acetonitrile, 0.1% formic acid) at 0.15–0.25 µg µL^−1^.

#### Serum sample preparation

To characterize proteins present in the serum matrix, proteins from 5 µL serum were precipitated by methanol crash (3:1 v/v ice-cold methanol, –20 °C, overnight). Pellets were collected by centrifugation and washed twice with 300 µL ice-cold methanol. The resulting protein pellets were processed using the PreOmics iST kit (PreOmics GmbH) according to the manufacturer’s instructions, including peptide purification and fractionation with the PreOmics fractionation add-on kit. Each sample was divided into three fractions prior to LC–MS/MS (“pre-fractionation”), yielding a total of 60 injections. Fractions were analyzed as independent runs but combined bioinformatically as a single sample. Purified peptides were dried under a nitrogen stream and resuspended in LC loading buffer (98:2 H₂O:acetonitrile, 0.1% formic acid) at 0.25 mg mL^−1^. Concentrations were determined using a NanoDrop spectrophotometer. For LC–MS/MS analysis, 1 µg peptide (4 µL injection volume) was loaded per fraction.

#### NanoLC–MS/MS analysis

All digested peptides were analyzed using an UltiMate 3000 RSLCnano system (ThermoFisher) coupled to a Q Exactive Plus Hybrid Orbitrap mass spectrometer (ThermoFisher) via nanoelectrospray ionization. Peptides were separated on a C18 column with a linear gradient of 4–35% acetonitrile (0.1% formic acid) over 135 min. MS1 spectra were acquired in positive mode at 70,000 resolution (m/z 200) with an AGC target of 2×10⁵ and a maximum injection time of 100 ms. Data-dependent acquisition was used to fragment the top 20 most abundant precursor ions (charge >1) with a 1.5 m/z isolation window, normalized collision energy of 30, fixed first mass of 140 m/z, and dynamic exclusion of 30 s. MS2 spectra were acquired at 17,500 resolution with an AGC target of 1×10⁵ and a maximum injection time of 60 ms.

#### Data analysis

All raw data were processed in FragPipe (v19.1) using default parameters for LFQ–MBR workflows.^80, 81^ To generate a reference database of serum proteins, fractionated serum samples were searched as single experiments per replicate. Variable modifications included oxidation (+15.995 Da, Met) and carbamylation (+42.025 Da, Lys), with carbamidomethylation (+57.025 Da, Cys) as a fixed modification. Searches were conducted against the *Homo sapiens* canonical + isoforms reference proteome (Uniprot UP000005640, accessed Dec 2024) with a precursor tolerance of ±20 ppm and allowance of up to three missed cleavages. Peptide-level FDR was controlled at 1%. Label-free quantification was performed using IonQuant^82^ with match-between-runs FDR set at 1% (top 350 runs). High-confidence proteins (>0.98 probability, >2 unique peptides, observed in 2/3 replicates) were designated as the “serum-specific” FASTA. Peptides isolated by in-gel digestion from replicate samples (n=3) were searched against the in-house serum-specific FASTA using FragPipe (v19.1) under the same parameters. Protein abundances were restricted to high-confidence identifications (>0.95 probability, >2 unique peptides, detected in at least 2 of 3 replicates). Abundances were log₂-transformed and median-normalized across replicates. For each protein, the average normalized log₂ abundance and standard error of the mean (SEM) were calculated from detected values only; missing values were not imputed. Experimental groups were compared to controls using these normalized abundance values. Protein molecular weights (kDa), isoelectric points (pI), and biological process classifications were determined using a custom Python analysis tool developed in-house, available at https://github.com/FitzkeeLab/proteomics_tool. Venn diagram for protein overlap analysis was generated using the InteractiVenn web tool (https://www.interactivenn.net/).^83^

### CD experiments

For circular dichroism (CD) experiments, protein solutions were prepared containing 0.2 mg mL^−1^ BSA, 0.2 mg mL^−1^ Tf, 0.1 mg mL^−1^ Fn, and 0.4 mg mL^−1^ IgG. These concentrations were selected to obtain a good CD signal-to-noise ratio. All the mixtures were prepared as described above and incubated with 100 nM AuNPs for at least 2 h. These samples were then transferred to a quartz demountable cuvette having a 0.2 mm path length. CD spectra were collected using a JASCO J-1500 CD spectrometer, and data analysis was done using SpectraManager software.

### Cellular uptake studies

RAW 264.7 macrophages and MDA-MB-231 triple-negative breast cancer cells were seeded in triplicate in 6-well plates at a density of 200,000 cells per well with media comprised of Dulbecco’s modified Eagle’s medium (DMEM) with 10% FBS and 1% antibiotic-antimycotic and allowed to adhere for 24 h. AuNPs were added to the media from a stock solution, resulting in a final concentration of 100 µg mL^−1^. The cells were treated with the AuNPs in the media for 6 h and then washed with DPBS. Cells were then trypsinized and collected into 1.5 mL centrifuge tubes. The cells were washed and centrifuged twice at 200 x g for 5 minutes, and the supernatant was discarded. The cells were then imaged using TEM and quantified for Au concentration using ICP-MS.

### TEM imaging

The cells were initially fixed with a primary fixative containing 2% paraformaldehyde, 2.5% glutaraldehyde, and 2 mM CaCl_2_ in 0.1 M sodium cacodylate buffer (pH 7.4) for 2 h at room temperature. After washing, the cells underwent secondary fixation with 1% osmium tetroxide for 1 h. Following additional washing, dehydration was carried out through a graded ethanol series, starting at 30% and increasing stepwise to 100% ethanol. The dehydrated cells were then infiltrated with 100% epoxy resin and polymerized at 60 °C over 1-2 days. Using an ultramicrotome, thin sections of the embedded cells were cut and collected on Formvar-coated copper grids. The grids were stained using 1% uranyl acetate and allowed to dry for 2 h in a desiccator before examination. Imaging was performed on a JEOL 2100 transmission electron microscope with an accelerating voltage of 200 kV to obtain high-resolution images of the cell sections.

### ICP-MS experiments for cells

Calibration standard solutions were prepared from a 999 mg L^−1^ gold standard for ICP (Sigma #38168) in the concentration range of 1 ppb to 1 ppm. Cells were digested overnight using 250 µL of aqua regia. The digested cells were diluted up to 10 mL and then analyzed by ICP-MS. A calibration curve was plotted for the standards, and Au content in each sample was calculated from the curve.

### Cytotoxicity studies

The viability of RAW 264.7 murine macrophages (ATCC #TIB-71) and MDA-MB-231 triple-negative breast cancer cells (Gentarget Inc. #SC057-Bsd) after treatment with the AuNPs and protein-coated AuNPs was determined using the CellTiter-Glo Luminescent Cell Viability assay. The cells were cultured in media comprised of Dulbecco’s modified Eagle medium (DMEM) (Gibco, #11965118) with 10% FBS and 1% antibiotic-antimycotic. The cells were seeded in triplicate in a 96-well plate at a density of 20,000 cells per well. The cells were allowed to adhere and proliferate for 24 h before replacing the media with new media containing AuNPs at a concentration of 100 µg mL^−1^. The cells were incubated with the new AuNP-containing media for 24 h, after which the wells were emptied and washed with DPBS (Ca^2+^/Mg^2+^ free) (Gibco, #14190359). The glow assay was subsequently performed to measure the luminescence of the cells using a BioTek microplate reader.

### In vivo pharmacokinetics and biodistribution

All animal procedures were conducted following the University of Mississippi Institutional Animal Care and Use Committee and National Institutes of Health (NIH) guidelines. Female BALB/c mice (4-6 weeks old) were quarantined for 7 days with 12-hour dark/light cycles and unrestricted access to standard pellet feed and water. 4T1 breast cancer cells (1×10^6^) were injected into the mammary fat pad of 4-6 week old immunocompetent mice, and tumors were grown to ~800 mm^3^.

For pharmacokinetics, mice were randomized into seven groups (n=3): control, AuNPs, PEGylated AuNPs, AuNPs+BSA+Fn, AuNPs+Tf+Fn, AuNPs+BSA+IgG, and AuNPs+BSA+Tf+Fn+IgG. Each group received 150 μL of AuNP formulation (10×10^12^ particles) via tail vein injection. Blood samples were collected under isoflurane anesthesia (5% induction, 2% maintenance) at 1, 2, 6, 12, and 24 hours post-injection. These blood samples were stored in K2 EDTA tubes and processed for gold content analysis. Each blood sample was digested by adding 500 μL of freshly prepared aqua regia and allowed to digest completely. The digested samples were then diluted to a final volume of 10 mL using Milli-Q water and filtered through 0.4 μm syringe filters. Au content in the filtered samples was quantified using ICP-MS. Gold concentration versus time data was fitted to a two-compartment model using GraphPad Prism software. Pharmacokinetic parameters, such as mean residence time (MRT) and area under the curve (AUC), were calculated from the fitted data to evaluate the circulation behavior of different AuNP formulations.

For biodistribution studies, mice were humanely sacrificed 24 h post-injection, and major organs (heart, liver, lungs, spleen, kidneys, and tumors) were carefully excised. Each organ was weighed immediately after collection and homogenized individually using an IKA 5G homogenizer under strict protocols to prevent cross-contamination. The homogenized tissue samples were digested by adding 2 mL of freshly prepared aqua regia and incubated overnight under appropriate safety conditions. Following complete digestion, samples were diluted to a final volume of 50 mL with Milli-Q water and filtered through 0.4 μm syringe filters to remove any tissue debris. Au content in the filtered samples was quantified using ICP-MS and normalized to the tissue weight, expressed as Au concentration per gram of organ tissue (ppm g^−1^). This normalization enabled direct comparison of nanoparticle accumulation across organs of different sizes and between treatment groups. Statistical analysis was performed to assess significant differences in organ distribution patterns among the various AuNP formulations, with particular emphasis on tumor accumulation.

### Statistical analysis and data availability

Unless otherwise stated, all measurements are reported as the average and the standard error of the mean for at least three independently prepared samples. When technical replicates are used instead, this is explicitly noted in the text and figure legends. Unless otherwise noted, comparisons were performed using one-way ANOVA with GraphPad Prism 10 software. Pairwise post-hoc testing was performed using Tukey’s multiple comparisons test, and statistical significance was reported at the *α* = 0.05 significance level. All raw data from this study have been submitted to Zenodo and are available at https://doi.org/10.5281/zenodo.17869720/.

## Supporting Information

The Supporting Information is available free of charge at …

## Author Contributions

The manuscript was written through the contributions of all authors. All authors have given approval to the final version of the manuscript.

## Notes

The authors declare no competing financial interests.

## Acknowledgments

This work was supported by the National Science Foundation under awards OIA 2414443, and CBET 2405018. Support for the MSU NMR Facility was provided by the National Science Foundation under awards CHE/MCB 2304919 and DBI 2215258. In vivo studies were supported in part by the American Cancer Society under grant RSG-21-114-01-MM and by the National Science Foundation under award OIA-2414442. Portions of this manuscript were assisted by the use of an AI language model for grammar and phrasing refinement. The authors verified the accuracy and take full responsibility for its content.

