## Supplemental Information for "Precision control of nanoparticle delivery with engineered biomimetic protein coronas"

### Supporting Figures

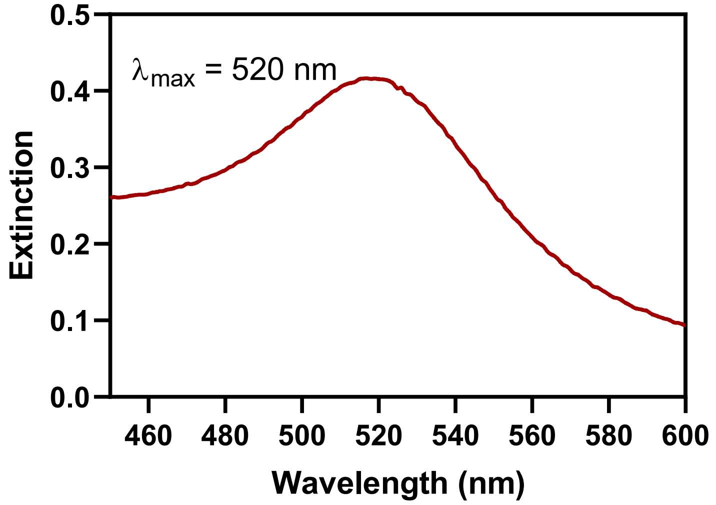

**Figure S1. UV spectra of synthesized 15 nm AuNPs.**

Data is shown for UV-Vis analysis of synthesized 15 nm AuNPs, and the $\lambda_{max}$ was found at 520 nm.

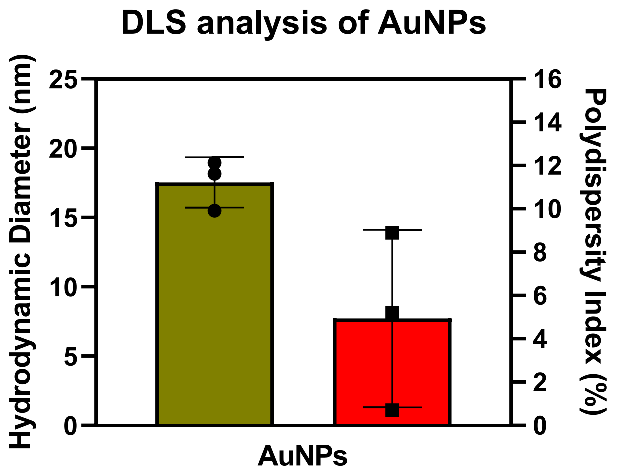

**Figure S2. Particle size distribution of synthesized AuNPs.**

The hydrodynamic diameter (brown, left) and polydispersity index (red, right) are shown for as-synthesized 15 nm AuNPs. Measurements were performed using an Anton Paar Dynamic Light Scattering (DLS) system at room temperature. Error bars represent the standard deviation from n = 3 independently prepared samples.

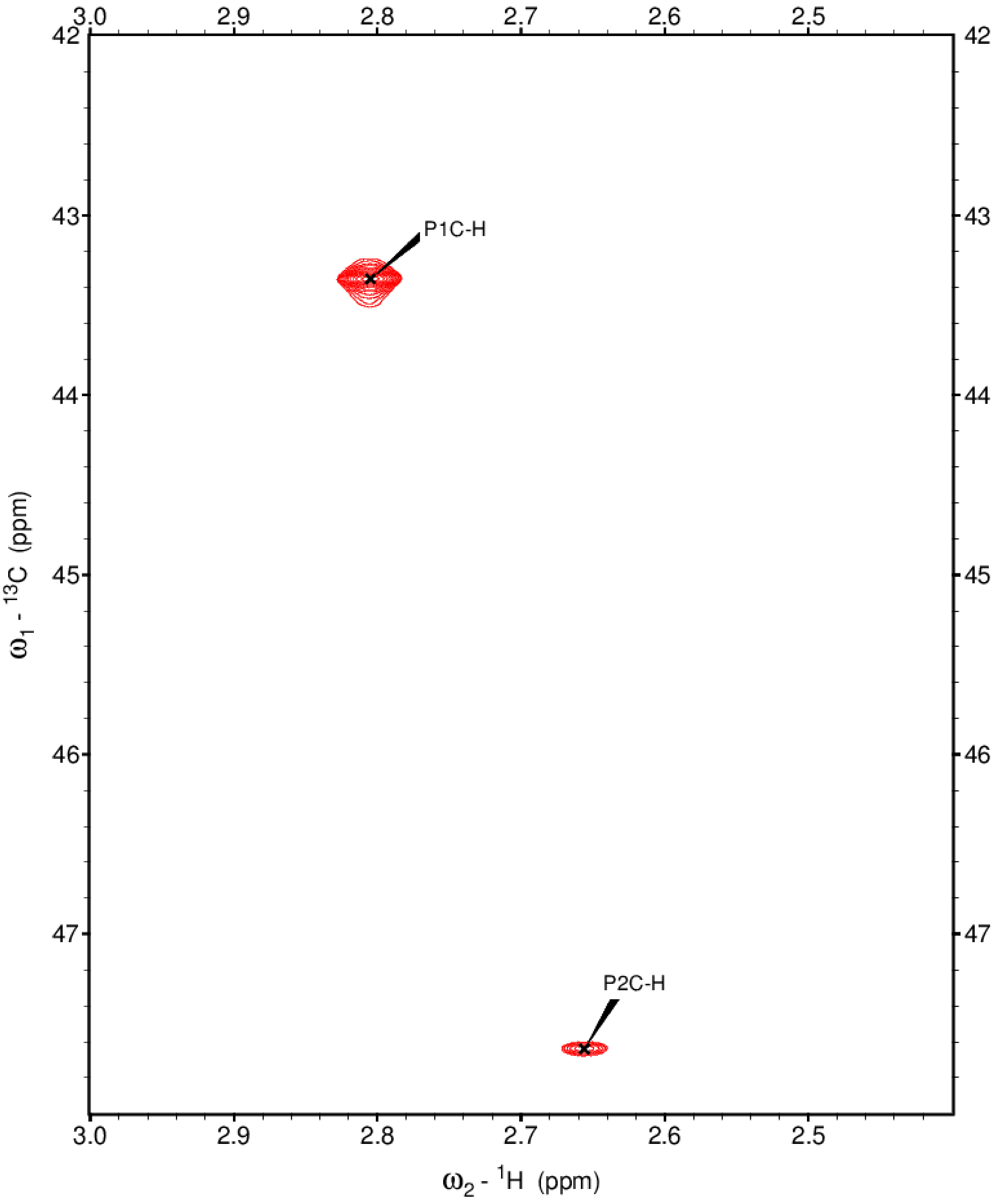

**Figure S3. NMR spectra of $^{13}C$ methylated proline as a quantitation standard**

$^{1}H$-$^{13}C$ HSQC NMR spectra of $^{13}C$ methylated proline, which is used as a quantitative standard for all the $^{13}C$ experiments. We observed two well-resolved peaks centered at 43.35 and 47.64 ppm in the $^{13}C$ dimension.

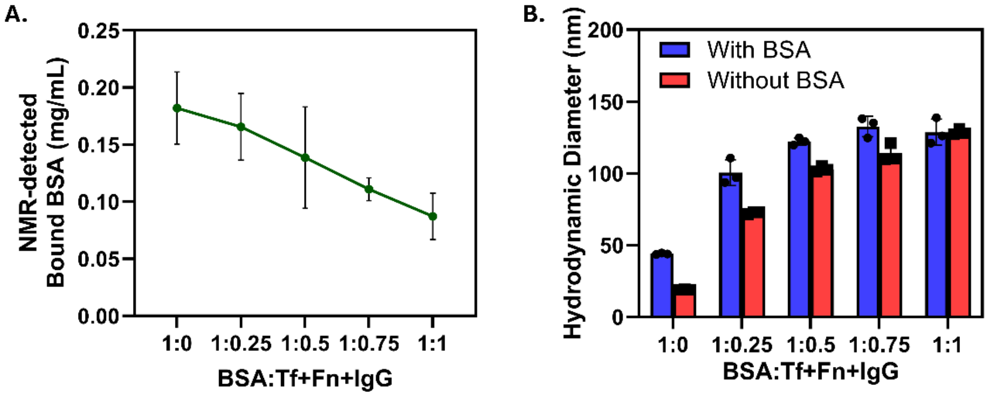

**Figure S4: Competitive interaction between proteins in a four-component mixture**

(A) NMR quantification showing decreasing bound BSA concentration as the total protein mixture concentration increases, demonstrating competitive displacement by other serum proteins (Tf, Fn, IgG) in the four-component system. (B) Dynamic light scattering measurements confirming protein corona formation with increasing mixture concentration, with hydrodynamic diameter reaching saturation at higher protein concentrations, indicating complete surface coverage.

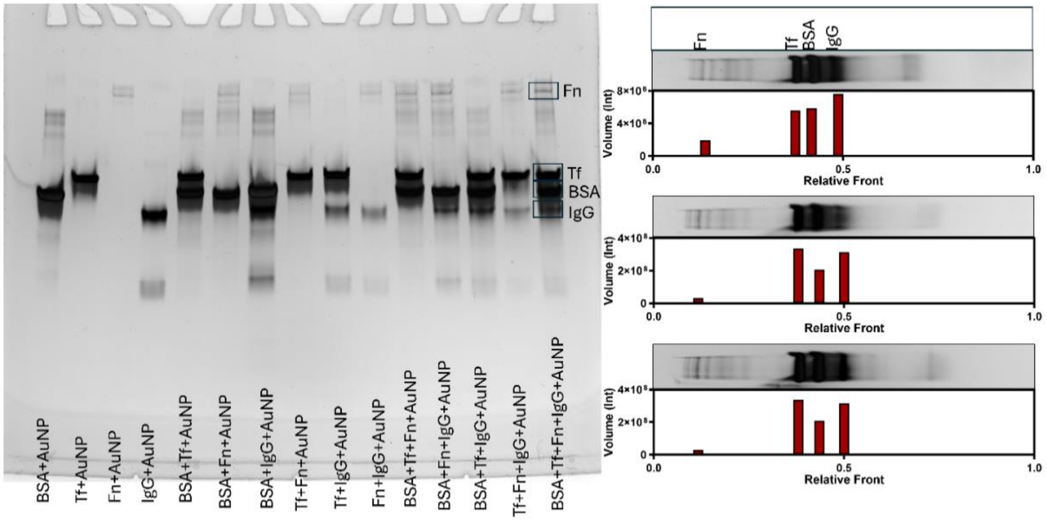

**Figure S5. SDS-PAGE of proteins bound to AuNPs**

The SDS-PAGE gel image displays the separation of protein mixtures bound onto gold nanoparticles (AuNPs). The presence of multiple bands in each lane indicates successful binding of various proteins to the AuNP surface. The differences in band intensities within each lane suggest variations in the binding affinities or amounts of the individual proteins present in the respective mixtures.

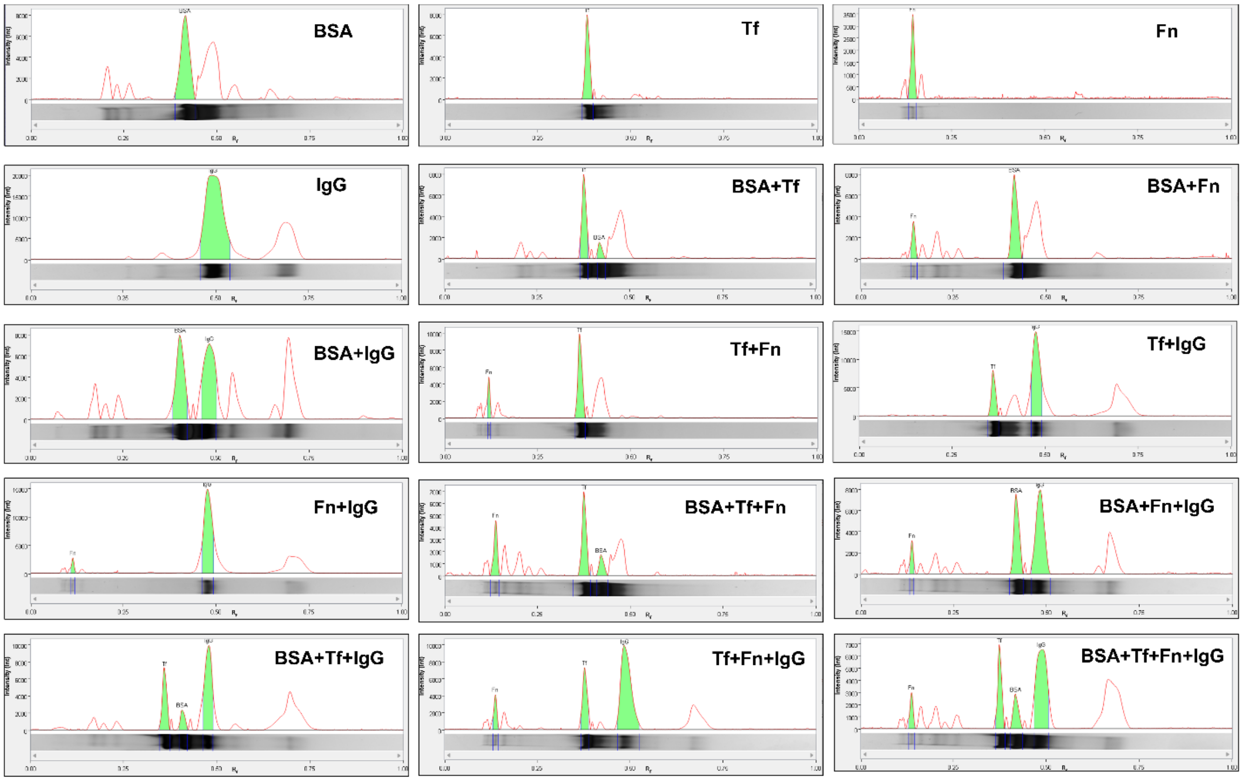

**Figure S6. Lane profiles of the gel densitometry data.**

This image presents the densitometric analysis of the SDS-PAGE gel lanes corresponding to the different protein mixtures bound onto AuNPs (Figure S5). Each graph represents the lane profile, which provides a quantitative representation of the band intensities within a particular lane. The x-axis of each graph corresponds to the migration distance or relative mobility of the proteins in the gel, while the y-axis represents the intensity of the bands. The presence of peaks in the lane profiles indicates the presence of individual protein bands, with the peak height reflecting the relative abundance of that particular protein to the AuNPs. This analysis was performed using Image Lab Software (Bio-Rad).

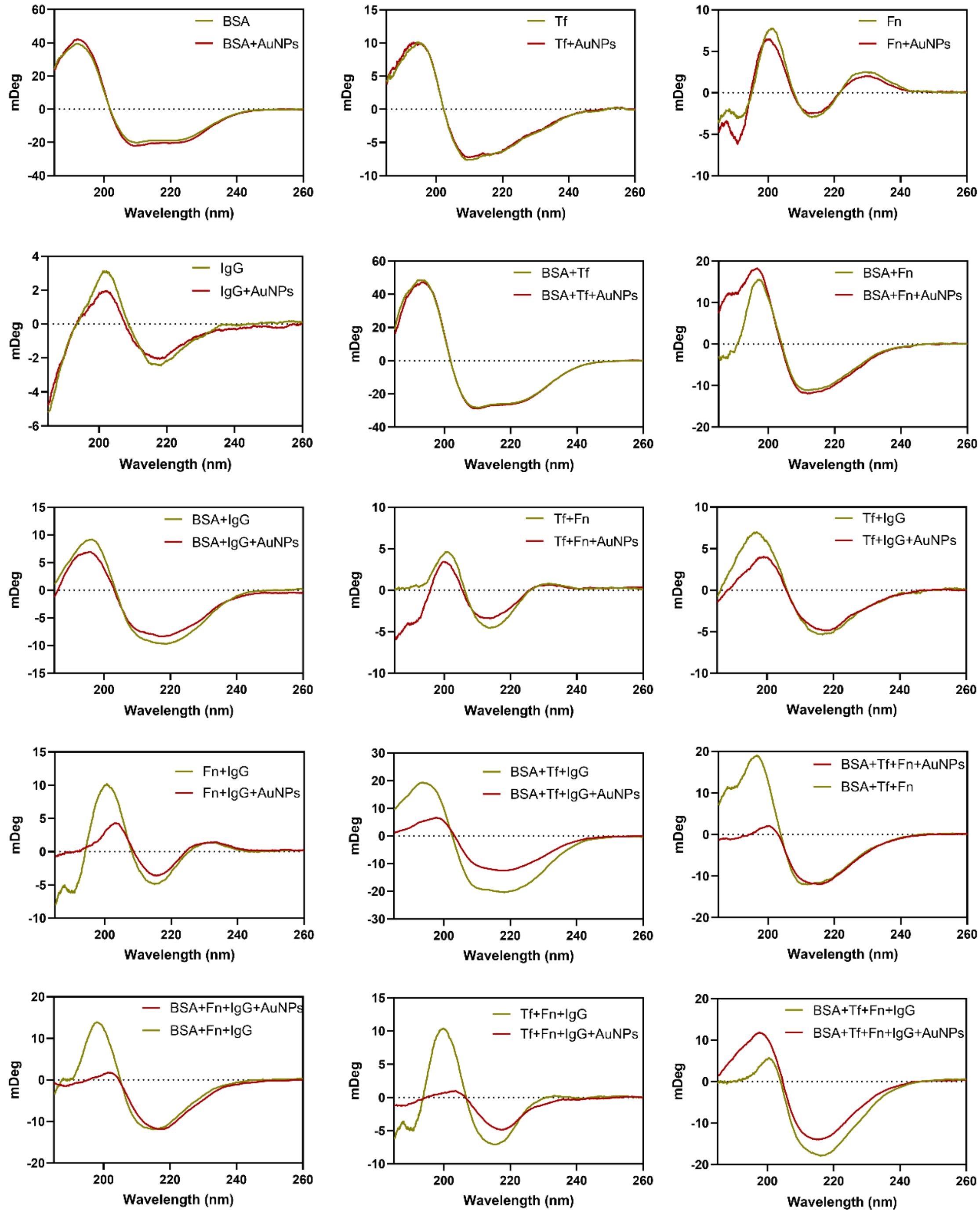

**Figure S7. CD spectra of all the mixtures**

CD spectra of protein mixtures with (red) and without (brown) AuNPs. The mixture composition is provided in the figure legend of each spectrum. We used 0.2 mg $mL^{-1}$ BSA, 0.2 mg $mL^{-1}$ Tf, 0.1 mg $mL^{-1}$ Fn, 0.4 mg $mL^{-1}$ IgG, and 100 nM AuNPs.

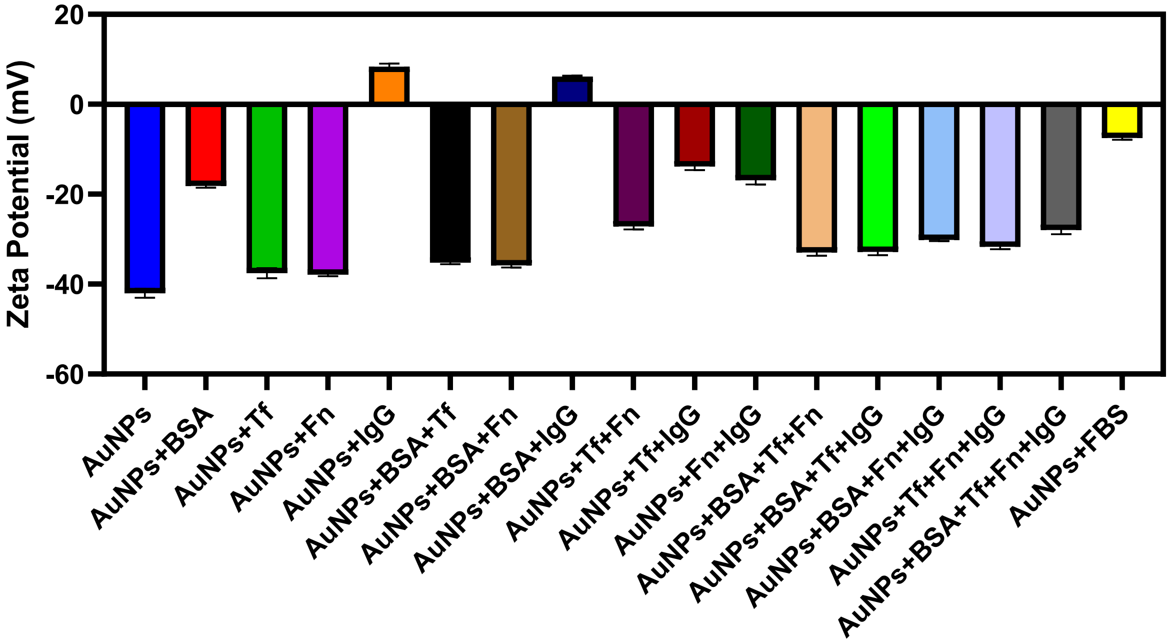

**Figure S8. Zeta Potential analysis of the mixtures**

Zeta potential analysis reveals how different protein adsorption affects the electrophoretic behavior of AuNPs. For example, a positive zeta potential of IgG became negative in the presence of Tf or Fn, indicating that Tf and Fn compete with IgG for binding to AuNPs. Error bars represent the standard error of the mean (SEM) of n = 3 repeated measurements on the same sample.

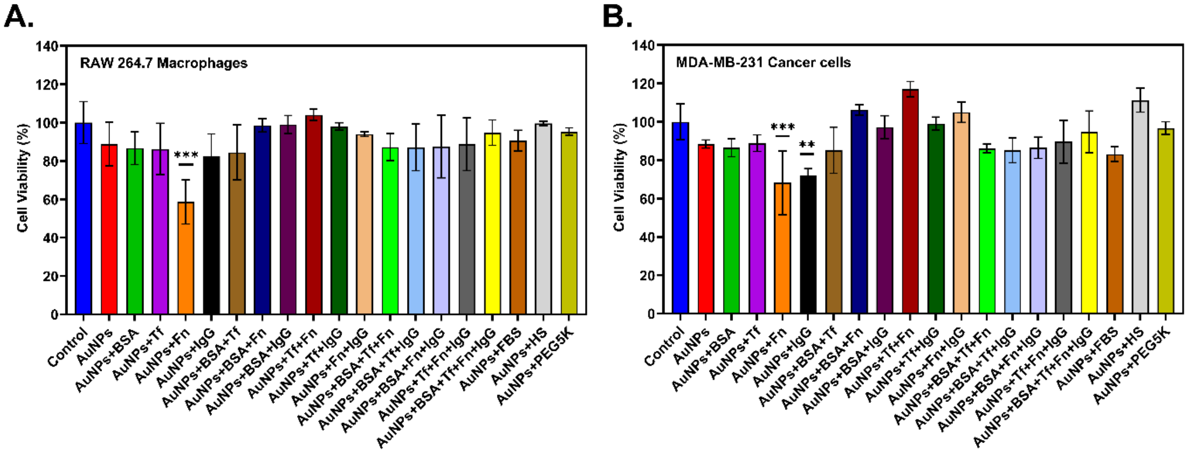

**Figure S9. Cytotoxicity assessment of different gold nanoparticle formulations in macrophages and breast cancer cells.**

(A) Cell viability (%) of RAW 264.7 macrophages after 24 h exposure to various gold nanoparticle formulations (100 μg $mL^{-1}$). (B) Cell viability (%) of MDA-MB-231 triple-negative breast cancer cells following 24 h treatment with different gold nanoparticle formulations (100 μg $mL^{-1}$). Data are presented as mean and standard deviation from three independent experiments.

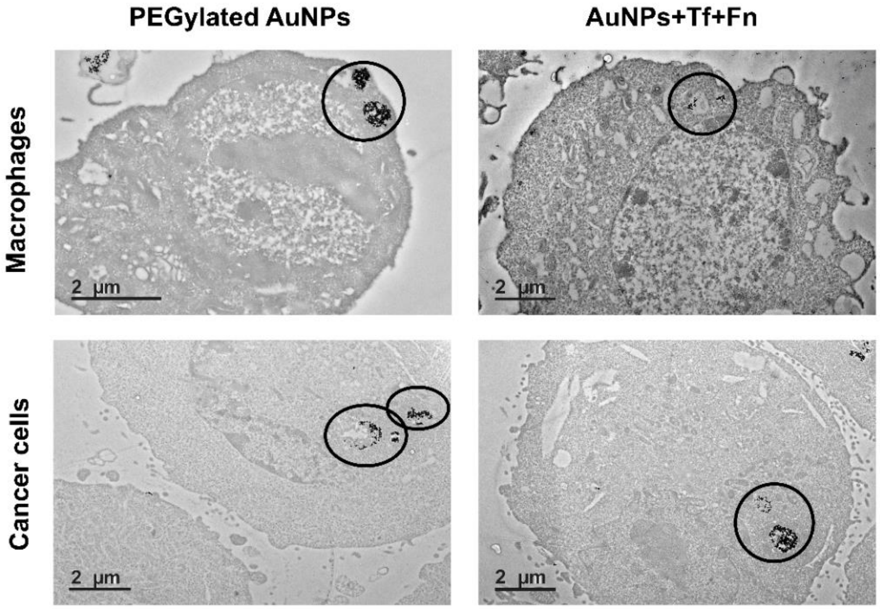

**Figure S10. TEM images of macrophages and cancer cells after incubation with PEGylated AuNPs, and Tf+Fn-coated AuNPs.**

TEM images showing the localization of gold nanoparticles (circled). Scale bars represent 2 μm.

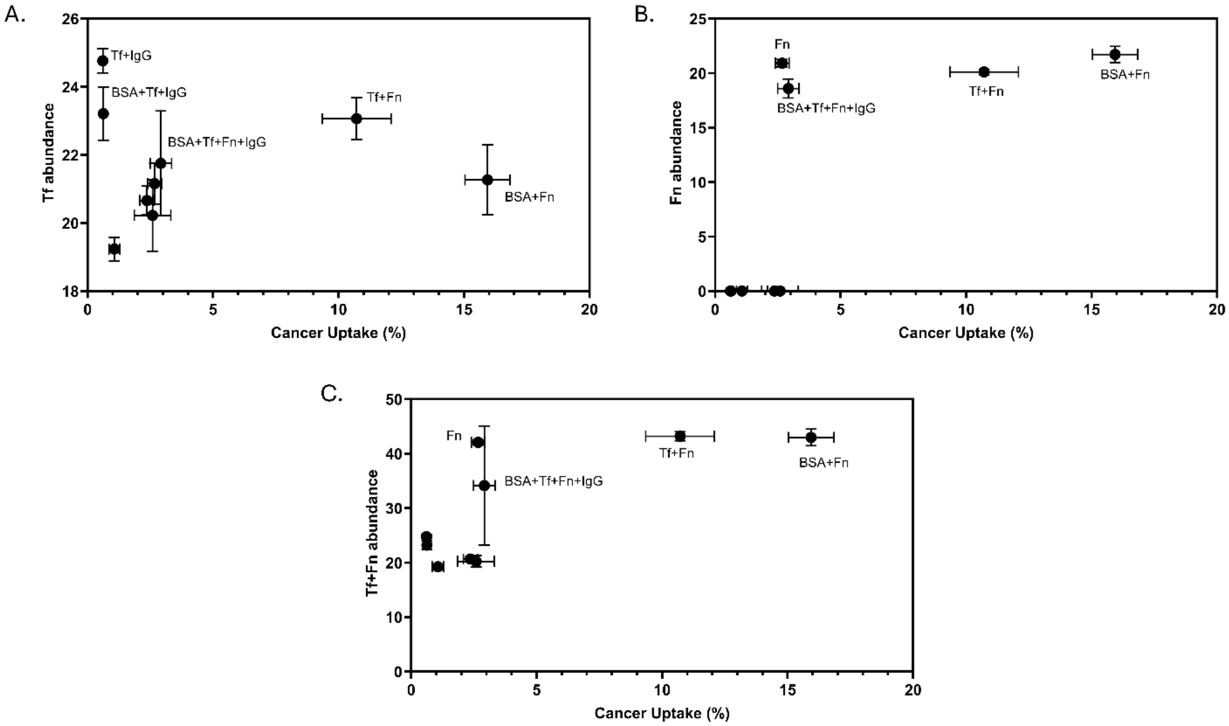

**Figure S11. Correlation between protein abundance in secondary coronas and cancer cell targeting efficiency.**

Scatter plots showing protein abundance (measured by LC-MS/MS after serum exposure) versus cancer cell uptake for (A) transferrin (Tf), (B) fibronectin (Fn), and (C) combined Tf+Fn abundance. The data reveal that cancer cell targeting is not simply determined by individual protein abundance, as some formulations with high Tf levels (panel A) show relatively low uptake. However, the combined Tf+Fn analysis (panel C) suggests that synergistic interactions between multiple targeting proteins may be more important than individual protein levels for achieving optimal cancer cell recognition.

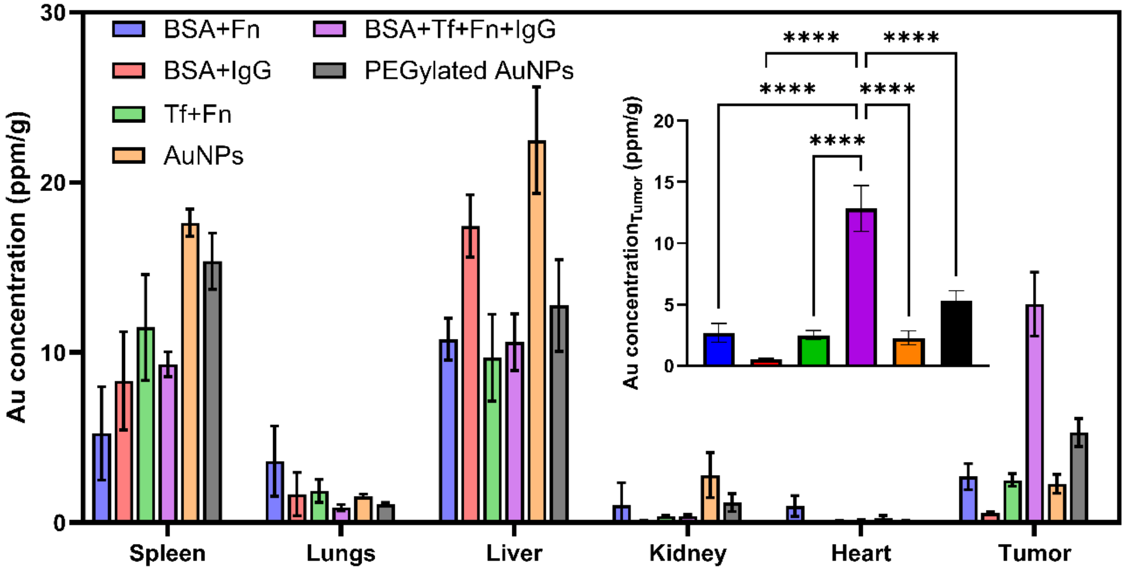

**Figure S12. Biodistribution analysis of different AuNP formulations in major organs and tumor tissue.**

The graph shows the biodistribution of various AuNP formulations (BSA+Fn, BSA+IgG, Tf+Fn, BSA+Tf+Fn+IgG, bare AuNPs, and PEGylated AuNPs) at 24 hours post-injection, expressed as gold concentration (ppm) per gram of tissue. The data reveals distinctive accumulation patterns across organs, with the BSA+Tf+Fn+IgG formulation demonstrating the highest tumor accumulation (~13 ppm/g) compared to all other formulations. Inset highlights tumor-specific accumulation showing statistically significant differences between formulations. Bare AuNPs show high accumulation in reticuloendothelial organs (spleen and liver), while engineered protein coronas modulate organ-specific distribution patterns. Error bars represent standard deviation (n=3). Statistical significance: ****$p < 0.0001$.

### Supporting Tables

#### Table S1. Average abundance of serum proteins in secondary coronas formed on differently functionalized AuNPs

Average abundance values ($log_2$-transformed LFQ intensities) of serum proteins in coronas formed on functionalized gold nanoparticles. Blank cells indicate protein absence in the respective corona.

| | | Average Abundance | | | | | | | | |
|---|---|---|---|---|---|---|---|---|---|---|
| **Protein ID** | **Description** | **AuNP** | **PEG5K** | **Fn** | **BSA +IgG** | **Tf+IgG** | **BSA+Tf +IgG** | **BSA +Fn** | **Tf+Fn** | **BSA+Tf+ Fn+IgG** |
| A0A075 B6I9 | Immunoglobulin lambda variable 7-46 | 18.16 | 19.76 | 19.71 | 18.36 | 17.29 | 19.26 | 19.97 | 17.69 | 19.51 |
| P01619 | Immunoglobulin kappa variable 3-20 | 20.89 | 19.92 | 21.35 | 19.91 | 20.11 | 21.17 | 21.53 | 19.77 | 20.04 |
| O14802 | DNA-directed RNA polymerase III subunit RPC1 | 21.50 | 22.22 | 18.56 | 18.26 | | | 22.70 | | |
| A0A075 B6H7 | Probable non-functional immunoglobulin kappa variable 3-7 | 21.14 | 20.87 | 21.66 | 20.19 | 20.81 | 21.54 | 22.18 | 20.63 | 21.02 |
| P01011 | Alpha-1-antichymotrypsin | | 20.83 | 19.93 | | | 18.26 | 18.99 | | 16.87 |
| A0A0A0 MRZ8 | Immunoglobulin kappa variable 3D-11 | 20.09 | 20.75 | 20.25 | 19.11 | 19.04 | 20.40 | 20.42 | 18.59 | 19.88 |
| P02663 | Alpha-S2-casein | | 21.12 | | | | | | | |
| A0A075 B6P5 | Immunoglobulin kappa variable 2-28 | 18.86 | 19.79 | 19.15 | | 19.00 | 18.64 | 19.14 | | 18.41 |
| P0DOY2 | Immunoglobulin lambda constant 2 | | | | | | 20.89 | 16.23 | | |
| A3KMH1 -2 | von Willebrand factor A domain-containing protein 8 | | 20.81 | 20.80 | 20.41 | 20.76 | 21.01 | 21.76 | 19.83 | 20.69 |
| P01591 | Immunoglobulin J chain | | 20.34 | 20.09 | | | | | | |
| O43866 | CD5 antigen-like | 16.53 | 19.40 | 16.60 | | | | | | |
| P13647 | Keratin, type II cytoskeletal 5 | 19.45 | 19.04 | 19.36 | 18.00 | | 16.55 | 19.37 | 18.43 | 15.49 |
| P02746 | Complement C1q subcomponent subunit B | 20.58 | 21.34 | 20.55 | 18.75 | | 18.79 | 19.46 | 17.04 | 17.76 |
| Q5T7N2 | LINE-1 type transposase domain-containing protein 1 | 24.00 | 23.82 | 23.83 | 22.20 | 22.50 | 22.73 | 24.52 | 22.34 | 23.26 |
| P01023 | Alpha-2-macroglobulin | 18.88 | 18.96 | 19.53 | 17.92 | 17.48 | 18.59 | 20.67 | 16.67 | 19.05 |
| P02768 | Albumin | 25.00 | 26.04 | 25.10 | 22.95 | 23.50 | 24.84 | 24.82 | 22.13 | 24.36 |
| P02769 | Albumin Bovine | 18.75 | | | 22.59 | 18.49 | 23.59 | 23.09 | 16.74 | 22.94 |
| P02743 | Serum amyloid P-component | | | | | | | 18.41 | | |
| P02647 | Apolipoprotein A-I | 18.27 | 19.16 | 18.22 | | | 18.29 | | | |
| P02656 | Apolipoprotein C-III | 19.86 | 19.14 | | | | 17.78 | | 16.60 | |
| P02747 | Complement C1q subcomponent subunit C | 21.05 | 21.14 | 21.60 | 19.32 | | 19.23 | 20.92 | 17.83 | 18.51 |

| | | | | | | | | | | |
|---|---|---|---|---|---|---|---|---|---|---|
| P09871 | Complement C1s subcomponent | 20.17 | 18.80 | 20.32 | 19.71 | | 19.35 | 21.17 | 16.63 | 18.52 |
| P01024 | Complement C3 | 23.61 | 21.27 | 22.91 | 20.48 | 19.33 | 21.14 | 22.74 | 19.72 | 21.37 |
| P02751-10 | Isoform 10 of Fibronectin | | | 20.91 | | | | 21.71 | 20.10 | 18.58 |
| P00738 | Haptoglobin | 21.68 | 21.35 | 21.72 | 19.71 | 19.13 | 20.03 | 21.75 | 18.42 | 19.34 |
| P01876-1 | Immunoglobulin heavy constant alpha 1 | 22.08 | 23.34 | 22.75 | 20.85 | 19.74 | 21.09 | 23.56 | 20.26 | 22.48 |
| P01857-1 | Immunoglobulin heavy constant gamma 1 | 25.90 | 26.13 | 26.21 | 25.01 | 26.50 | 26.05 | 26.75 | 25.06 | 25.94 |
| P01859-1 | Immunoglobulin heavy constant gamma 2 | 23.16 | 23.51 | 23.45 | 22.09 | 23.15 | 22.72 | 23.97 | 21.58 | 21.75 |
| P01871-1 | Immunoglobulin heavy constant mu | 22.97 | 23.24 | 23.50 | 21.16 | 21.25 | 22.17 | 22.95 | 20.86 | 22.22 |
| P01834 | Immunoglobulin kappa constant | 21.44 | 21.88 | 21.66 | 19.65 | 21.57 | 21.21 | 22.18 | 19.05 | 20.51 |
| P04264 | Keratin, type II cytoskeletal 1 | 23.06 | 22.99 | 23.77 | 21.57 | 22.24 | 20.90 | 23.10 | 21.80 | 21.17 |
| P13645 | Keratin, type I cytoskeletal 10 | 22.78 | 22.46 | 23.21 | 19.49 | 21.80 | 20.31 | 22.80 | 21.21 | 20.49 |
| P02533 | Keratin, type I cytoskeletal 14 | 19.85 | 19.38 | 19.80 | 17.98 | 17.57 | 17.41 | 19.30 | 18.92 | 17.82 |
| P35527 | Keratin, type I cytoskeletal 9 | 21.33 | 21.16 | 22.57 | 21.03 | 21.56 | 19.87 | 22.06 | 20.39 | 19.73 |
| P01009 | Alpha-1-antitrypsin | | 20.04 | 20.65 | 18.61 | 19.11 | 19.83 | 20.48 | 17.29 | 18.77 |
| P02787 | Serotransferrin | 20.66 | 20.22 | 21.16 | 19.24 | 24.76 | 23.21 | 21.27 | 23.06 | 21.75 |

**Table S2. Normalized protein abundance ratios relative to bare AuNP**

Values represent protein abundance normalized to the AuNP corona, where values >1 indicate higher abundance and values <1 indicate lower abundance compared to bare AuNP. Blank cells indicate the protein was not detected in the corresponding functionalized nanoparticle corona, while # indicates the protein was absent in the bare AuNP sample, preventing ratio calculation.

| | | Ratio Compared to AuNP | | | | | | | |
|---|---|---|---|---|---|---|---|---|---|
| | **Description** | **PEG5K** | **Fn** | **BSA +IgG** | **Tf+IgG** | **BSA+Tf +IgG** | **BSA+Fn** | **Tf+Fn** | **BSA+Tf+ Fn+IgG** |
| A0A075B6I9 | Immunoglobulin lambda variable 7-46 | 1.08 | 1.08 | 1.01 | 0.95 | 1.06 | 1.09 | 0.97 | 1.07 |
| P01619 | Immunoglobulin kappa variable 3-20 | 0.95 | 1.02 | 0.95 | 0.96 | 1.01 | 1.03 | 0.94 | 0.95 |
| O14802 | DNA-directed RNA polymerase III subunit RPC1 | 1.03 | 0.86 | 0.85 | | | 1.05 | | |
| A0A075B6H7 | Probable non-functional immunoglobulin kappa variable 3-7 | 0.98 | 1.02 | 0.95 | 0.98 | 1.01 | 1.04 | 0.97 | 0.99 |
| P01011 | Alpha-1-antichymotrypsin | # | # | | | # | # | | # |
| A0A0A0MRZ8 | Immunoglobulin kappa variable 3D-11 | 1.03 | 1.00 | 0.95 | 0.94 | 1.01 | 1.01 | 0.92 | 0.98 |
| P02663 | Alpha-S2-casein | # | | | | | | | |
| A0A075B6P5 | Immunoglobulin kappa variable 2-28 | 1.04 | 1.01 | | 1.00 | 0.98 | 1.01 | | 0.97 |
| P0DOY2 | Immunoglobulin lambda constant 2 | | | | | # | # | | |
| A3KMH1-2 | von Willebrand factor A domain-containing protein 8 | # | # | # | # | # | # | # | # |
| P01591 | Immunoglobulin J chain | # | # | | | | | | |
| O43866 | CD5 antigen-like | 1.17 | 1.00 | | | | | | |
| P13647 | Keratin, type II cytoskeletal 5 | 0.97 | 0.99 | 0.92 | | 0.85 | 0.99 | 0.94 | 0.79 |
| P02746 | Complement C1q subcomponent subunit B | 1.03 | 0.99 | 0.91 | | 0.91 | 0.94 | 0.82 | 0.86 |
| Q5T7N2 | LINE-1 type transposase domain-containing protein 1 | 0.99 | 0.99 | 0.92 | 0.93 | 0.94 | 1.02 | 0.93 | 0.96 |
| P01023 | Alpha-2-macroglobulin | 1.00 | 1.03 | 0.94 | 0.92 | 0.98 | 1.09 | 0.88 | 1.00 |
| P02768 | Albumin | 1.04 | 1.00 | 0.91 | 0.94 | 0.99 | 0.99 | 0.88 | 0.97 |
| P02769 | Albumin_Bovine | | | 1.20 | 0.98 | 1.25 | 1.23 | 0.89 | 1.22 |
| P02743 | Serum amyloid P-component | | | | | | # | | |
| P02647 | Apolipoprotein A-I | 1.04 | 0.99 | | | 1.00 | | | |
| P02656 | Apolipoprotein C-III | 0.96 | | | | 0.89 | | 0.83 | |
| P02747 | Complement C1q subcomponent subunit C | 1.00 | 1.02 | 0.91 | | 0.91 | 0.99 | 0.84 | 0.87 |

| | | | | | | | | | |
|---|---|---|---|---|---|---|---|---|---|
| P09871 | Complement C1s subcomponent | 0.93 | 1.00 | 0.97 | | 0.95 | 1.04 | 0.82 | 0.91 |
| P01024 | Complement C3 | 0.90 | 0.97 | 0.86 | 0.81 | 0.89 | 0.96 | 0.83 | 0.90 |
| P02751-10 | Isoform 10 of Fibronectin | | # | | | | # | # | # |
| P00738 | Haptoglobin | 0.98 | 1.00 | 0.90 | 0.88 | 0.92 | 1.00 | 0.84 | 0.89 |
| P01876-1 | Immunoglobulin heavy constant alpha 1 | 1.05 | 1.03 | 0.94 | 0.89 | 0.95 | 1.06 | 0.91 | 1.01 |
| P01857-1 | Immunoglobulin heavy constant gamma 1 | 1.00 | 1.01 | 0.96 | 1.02 | 1.00 | 1.03 | 0.96 | 1.00 |
| P01859-1 | Immunoglobulin heavy constant gamma 2 | 1.01 | 1.01 | 0.95 | 0.99 | 0.98 | 1.03 | 0.93 | 0.93 |
| P01871-1 | Immunoglobulin heavy constant mu | 1.01 | 1.02 | 0.92 | 0.92 | 0.96 | 0.99 | 0.90 | 0.96 |
| P01834 | Immunoglobulin kappa constant | 1.02 | 1.01 | 0.91 | 1.00 | 0.98 | 1.03 | 0.88 | 0.95 |
| P04264 | Keratin, type II cytoskeletal 1 | 0.99 | 1.03 | 0.93 | 0.96 | 0.90 | 1.00 | 0.94 | 0.91 |
| P13645 | Keratin, type I cytoskeletal 10 | 0.98 | 1.01 | 0.85 | 0.95 | 0.89 | 1.00 | 0.93 | 0.89 |
| P02533 | Keratin, type I cytoskeletal 14 | 0.97 | 0.99 | 0.90 | 0.88 | 0.87 | 0.97 | 0.95 | 0.89 |
| P35527 | Keratin, type I cytoskeletal 9 | 0.99 | 1.05 | 0.98 | 1.01 | 0.93 | 1.03 | 0.95 | 0.92 |
| P01009 | Alpha-1-antitrypsin | # | # | # | # | # | # | # | # |
| P02787 | Serotransferrin | 0.97 | 1.02 | 0.93 | 1.19 | 1.12 | 1.02 | 1.11 | 1.05 |
